# Inhibitory TIGIT signalling is dependent on T cell receptor activation

**DOI:** 10.1101/2025.05.08.652881

**Authors:** William H. Zammit, Lucy Le Maistre, Thomas A.E. Elliot, Stuart A. Cain, Martin J. Humphries, Daniel M. Davis, Jonathan D. Worboys

## Abstract

TIGIT is an immune checkpoint receptor that can signal via cytoplasmic ITT-like and ITIM motifs to regulate T cell function. The signalling molecules that mediate inhibitory TIGIT signalling in T cells remain poorly defined, and it is not clear how TIGIT activation is regulated. Here, proximity proteomics was employed in Jurkat T cells to identify TIGIT-associating proteins upon engagement with its ligand CD155. This identified several ligation-specific TIGIT interactors, including proteins involved in signalling (Grb2 and SOS1), cytoskeletal regulation (CD2AP and SdcBP), and endocytosis (IST1 and SNX3). A TIGIT mutant (Y225A/Y231A) incapable of signalling via its inhibitory motifs prevented the recruitment of these proteins, indicating inhibitory signalling-specific engagement of these pathways. Strikingly, T cell receptor (TCR) stimulation was also needed for TIGIT to engage with these pathways. Mechanistically, phosphorylation of TIGIT required both CD155 ligation and TCR activation, which resulted in signalling and internalisation. These findings demonstrate that TIGIT signalling is context-dependent and restricted to T cells receiving simultaneous TCR activation. This establishes a regulatory mechanism that limits checkpoint control to when functionally required.

## Main

T-cell immunoreceptor with Ig and ITIM domains (TIGIT) is an immune checkpoint receptor targeted in a range of phase III oncological clinical trials^1^. TIGIT binds to nectin and nectin-like molecules on antigen-presenting cells (APC) and cancer cells, interacting with CD155 with the highest affinity^2–4^. TIGIT can inhibit T cells directly through its signalling^5–8^ and by impairing the signalling of the co-stimulatory molecule CD226, either by competing for CD155 binding to CD226 *in trans*^2–4^, or by interacting with CD226 *in cis* to prevent it from forming homodimers^6,7^. In pre-clinical tumour models, TIGIT blockade is most effective as a combination therapy with PD-1 or PD-L1 blockade^5,6,9,10^, for reasons that are not clear. One hypothesis is that inhibition from TIGIT and PD-1 converges within cells to prevent CD226 signalling and only blocking both TIGIT and PD-1 enables complete CD226-driven co-stimulation^7^. However, tumour-infiltrating lymphocytes rarely co-express both TIGIT and CD226^11^ and, since TIGIT blockade disrupts both TIGIT-CD155 and TIGIT-CD226 interactions, it remains unclear which pathway predominantly drives inhibition in tumours. In contrast, distinct T cell states can be observed following TIGIT or PD-L1 monotherapy in murine tumour models, which may indicate their co-blockade could be synergistic through non-convergent mechanisms^8^.

Beyond its ability to curtail CD226 signalling, TIGIT function in T cells is poorly understood. TIGIT contains inhibitory ITT-like and ITIM motifs in its cytoplasmic tail^2–4^, which are essential for NK cell inhibition^12,13^. In NK cells, phosphorylation of the tyrosine residues within the ITT-like domain and the ITIM (Y225 and Y231 in human sequence, respectively) enables binding of the adaptor proteins Grb2 or β-arrestin. These can then recruit the inhibitory phosphatase SHIP-1 to inhibit downstream MAPK, Akt, and NFκB signalling to supress cytolytic cell function and inflammatory cytokine production^13,14^. In T cells, the role of inhibitory signalling has been debated, and it has been difficult to elucidate whether the same molecules transduce TIGIT signals. For example, a recent study was unable to precipitate Grb2 with TIGIT in T cells following ligation, although the interaction could be observed with a phosphorylated peptide pull down in cell lysates^7^. Functionally, TIGIT has demonstrated intrinsic signalling in both primary T cells^15^ and in an inhibitory motif-dependent manner in the Jurkat T cell line^11^. TIGIT ligation causes it to cluster to within nanometres of T cell receptor (TCR) clusters and it has been proposed that this nano-proximity is important for its inhibitory activity^11,16^. Thus, despite the importance of TIGIT as an immunoregulator in T cells and its significant therapeutic potential, it is still unclear how TIGIT signalling inhibits T cells.

Here, we employed proximity proteomics to map TIGIT signalling associations in model human T cells in an unbiased manner. TIGIT interactome maps were generated under different conditions to discern ligation-, TCR activation-, and inhibitory signalling-dependent interactions. This identified a range of previously undescribed associating proteins, including the cytoskeletal and membrane adaptor proteins CD2-associated protein (CD2AP), syntenin-1 (SdcBP), and the ESCRT protein IST1. Unexpectedly, ligation-dependent inhibitory signalling associations required T cell activation. Mechanistically TIGIT phosphorylation following ligation only occurred in activated T cells, leading to its signalling and internalisation. This provides evidence that the co-clustering of ligated TIGIT and TCR provides the stimulation necessary for TIGIT activity in T cells, establishing a spatially defined regulation of checkpoint function.

### Generation and functional validation of TIGIT-APEX2 fusion proteins

As inhibitory TIGIT signalling in T cells is poorly defined, we employed a proximity proteomic approach to identify signalling molecules that associate with TIGIT upon its activation. TIGIT has a single extracellular immunoglobulin domain that binds its ligands, a single-pass transmembrane domain, and a short cytoplasmic domain containing the ITT-like and ITIM signalling motifs^2,4^. Thus, lentiviral TIGIT constructs were designed containing the engineered peroxidase APEX2^17^ at the cytoplasmic tail to identify intracellular associating proteins (Fig. 1a). APEX2 is an engineered peroxidase that catalyses the production of reactive biotin-phenoxyl radicals in the presence of biotin-phenol (BP) and H_2_O_2_, and thereby biotinylate proximal proteins. TIGIT was fused to APEX2 with either a GSW linker (TIGIT-GSW-APEX2; GSW) or a self-cleaving T2A peptide (TIGIT-T2A-APEX2; T2A), which provides an APEX2 cytoplasmic control. Jurkat cells were transduced with TIGIT-APEX2 constructs, and expression was confirmed by both flow cytometry (Fig. 1b) and western blotting (Fig. 1c, top panel). The functionality of TIGIT-APEX2 was tested by analysing protein biotinylation patterns in the presence or absence of BP and H_2_O_2_. In both GSW and T2A cells, a range of proteins bound streptavidin in a BP- and H_2_O_2_-dependent manner (Fig. 1c, bottom panel), demonstrating efficient protein biotinylation.

**Figure 1.**
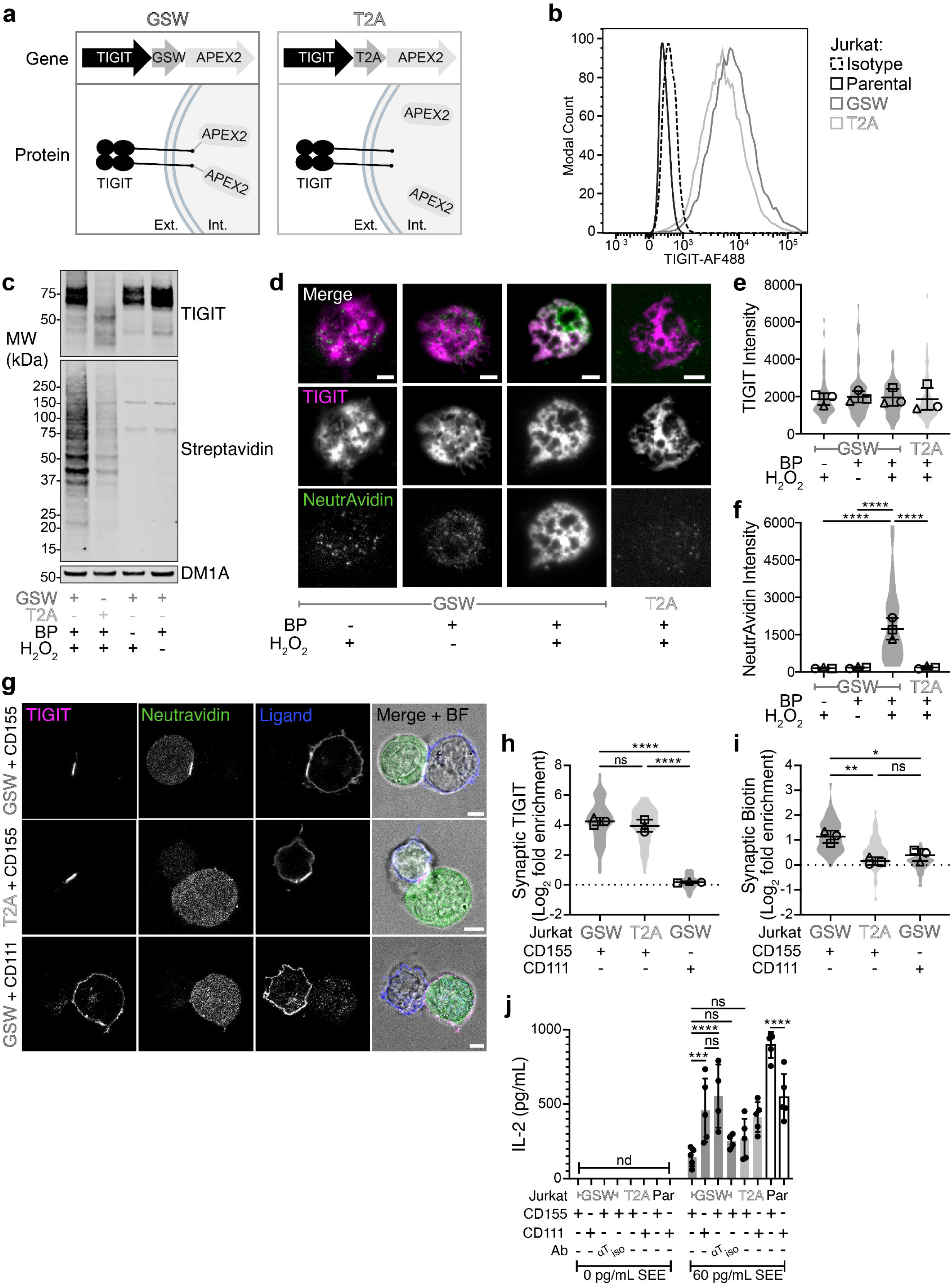
Generation of a functional TIGIT-APEX2 fusion protein to perform proximity proteomics in T cells. **a.** Schematic depicting the expression of TIGIT and APEX2 in Jurkat T cells. TIGIT is expressed fused to APEX2 with either a non-cleavable GSW linker or a cleavable T2A linker, as shown in the corresponding gene and protein structures. Ext = Extracellular, Int = Intracellular. **b.** Flow cytometric analysis showing the expression of TIGIT in TIGIT-GSW-APEX2 (GSW, dark grey), TIGIT-T2A-APEX2 (T2A, light grey), and parental Jurkat T cells (black) together with an isotype-matched control (dashed black). **c.** Western blot showing TIGIT and biotinylated proteins in GSW and T2A Jurkat cells preincubated with biotin-phenol (BP, 1mM, 30 min) and/or H_2_O_2_ (0.25 mM, 2 min). Controls without BP or H_2_O_2_ are included for GSW cells. A loading control blot for α-Tubulin (DM1A) is also shown below from the same lysates. Molecular weight markers are depicted on the left. **d.** Representative Total Internal Reflection Fluorescence (TIRF) microscopy images of TIGIT (magenta) at the immune synapse of GSW and T2A Jurkat cells that have interacted with Planar Lipid Bilayers (PLB) loaded with ICAM-1 (100 molecules/μm^2^) and CD155 (400 molecules/μm^2^) for 20 mins. Cells were preincubated with BP and H_2_O_2_ was added to the cells after 20 mins of interaction with PLBs, where indicated. NeutrAvidin (green) is used to mark biotinylated proteins. Controls without BP or H_2_O_2_ are included for GSW cells. A merged fluorescence-BF image is also provided. Scale bars = 5 μm. **e, f.** Mean pixel intensity values of TIGIT (**e**) and NeutrAvidin (**f**) in Jurkat cells from the TIRF imaging shown in **d** (±S.D.; n=3 independent experiments; significant adjusted *P* values from a one-way ANOVA with Tukey’s multiple comparisons are shown; violin plots display individual cell variability). **g.** Confocal microscopy images showing TIGIT (magenta) in GSW and T2A Jurkat cells that were conjugated with different Raji cell populations (expressing either CD111 or CD155) for 5 mins, as indicated on the left of the panel. Jurkat cells were preincubated with BP, and H_2_O_2_ was added after conjugation with Raji cells. NeutrAvidin (green) is used to mark biotinylated proteins and a V5 stain labels expressed ligands on Raji cells (blue). A merged fluorescence-BF image is provided. Scale bars = 5 μm. **h,i.** Mean log_2_ fold change in synaptic TIGIT (**h**) and NeutrAvidin (**i**) enrichment in Jurkat cells, from the conjugates shown in **g** (±S.D.; n=3 independent experiments; adjusted *P* values from a one-way ANOVA with Tukey’s multiple comparisons are given; ns = not significant; violin plots display individual cell variability). **j.** ELISA data showing the amount of IL-2 released from either parental or different forms of TIGIT-APEX2 expressing Jurkat cells after co-incubation with SEE-pulsed Raji cells for 6 h. GSW cells were also pre-incubated with an antagonistic TIGIT antibody (αT) or an isotype-matched control (iso), as indicated (±S.D.; n=4-5 independent experiments; adjusted *P* values from a one-way ANOVA with Tukey’s multiple comparisons are given; ns = not significant; nd = not detected).

To further validate functionality, the ability of TIGIT-APEX2 to bind CD155 was tested. Previously, we demonstrated that TIGIT clusters at the immune synapse (IS) upon engagement with CD155^11^. Thus, the distribution of both TIGIT and the resulting biotinylated proteins in Jurkat GSW and T2A cells upon ligation was visualised. Initially, planar lipid bilayers (PLB) containing the adhesion molecule ICAM-1 and CD155 to facilitate cell binding and TIGIT clustering, respectively, were generated. Both GSW and T2A cells were added to PLBs, proximity labelling experiments performed, before cells were fixed and labelled with a fluorescent TIGIT antibody and NeutrAvidin to visualise TIGIT and the resulting biotinylated proteins, respectively. Synaptic molecules were imaged using Total Internal Reflection Fluorescence (TIRF) microscopy (Fig. 1d). TIGIT clustered at the IS in all conditions, as expected. However, a significant enrichment of biotinylated proteins at the IS was only observed in GSW cells pre-incubated with both BP and H_2_O_2_ (Fig. 1d-f). Next, the synaptic enrichment of TIGIT and biotinylation was tested by imaging conjugates of TIGIT-APEX2-expressing Jurkat cells and Raji cells expressing either CD155 or a control non-binding ligand, CD111, with confocal microscopy (Fig. 1g). Both TIGIT-APEX2 proteins were enriched at the IS of Jurkat-Raji conjugates when Raji cells expressed CD155 (Mean Log_2_ fold-enrichment of ∼4) but not when they expressed CD111 (Mean Log_2_ fold-change of ∼0), as expected (Fig. 1g, h). However, biotinylated proteins only accumulated at the IS when GSW cells were conjugated with Raji cells expressing CD155 (Mean Log_2_ fold-enrichment of ∼1, compared to ∼0.1 with Raji-CD111; Fig. 1g, i). Taken together, these findings demonstrate that TIGIT-APEX2 can bind to CD155, localise and cluster at the IS and biotinylate proximal proteins.

Finally, the inhibitory signalling potential of TIGIT-APEX2 proteins was tested. We stimulated Jurkat GSW and T2A cells with Raji-CD111 or CD155 cells pulsed with the superantigen Staphylococcal Enterotoxin E (SEE). Both GSW and T2A cells co-cultured with SEE-pulsed Raji-CD155 cells released less IL-2 compared to when co-cultured with SEE-pulsed Raji-CD111 cells (∼70% and ∼40% less, respectively; Fig. 1j). The reduction in IL-2 released from GSW Jurkat cells conjugated with Raji-CD155 was restored with an antagonistic TIGIT antibody, but not an isotype-matched control. Conversely, parental Jurkat cells released more IL-2 when co-cultured with Raji-CD155 cells than with Raji-CD111 cells, likely due to low levels of CD226 expressed on Jurkat cells^11^. Thus, fusion of APEX2 to TIGIT did not impair its ability to inhibit T cell activation. Collectively, these results validate the APEX2 platform to investigate inhibitory TIGIT signalling in T cells.

### TIGIT engages signalling, cytoskeletal and endocytosis pathways upon ligation

Initially, we sought to elucidate the signalling molecules engaged by TIGIT upon CD155 ligation. Thus, the differential proximity proteomes of GSW in Jurkat cells conjugated with SEE-pulsed Raji-CD155 compared to Raji-CD111 was analysed (Fig. 2a). Proximity labelling in GSW cells was performed following a 5-minute incubation with the different SEE-pulsed Raji cells, resulting in protein biotinylation patterns that were indistinguishable by SDS-PAGE (Fig. 2b, Supplementary fig.1a, b). Label-free quantification of proteins by mass spectrometry measured approximately 3,500 proteins from each affinity-purified eluate from each conjugate condition and the T2A background control (Supplementary Fig. 1c, see methods for further experimental details). Principal component analysis (PCA) of the quantified proteins revealed a distinct separation between each of the conditions (Fig. 2c). The first principal component (PC1), accounted for 41.6% of the variability, and separated the conjugated GSW proximity proteomes from the control proteomes (T2A). The second principal component (PC2), accounted for 14.7% of the total variability, and separated the GSW proximity proteomes according to their ligation. Pairwise comparisons indicated a high similarity within the replicates for each condition with a strong Pearson correlation score between the GSW ligation conditions (Fig. 2d). As with the PCA, this highlights more similarity between the conjugated conditions than the background control. To compare the TIGIT interactome between different ligation conditions, we initially filtered both GSW proteomes using the background T2A control interactome, to remove non-specific interactors (see methods for further details). Differential enrichment of proteins within the CD155-ligated proximity proteome as compared to the CD111 control revealed 26 proteins more proximal to TIGIT upon CD155 ligation, and 11 proteins more proximal in the CD111 control (Fig. 2e, f). Grb2, an adaptor protein previously established as a TIGIT interactor in NK cells^7,13^, was more proximal upon CD155-ligation, providing validation that this approach can identify bone fide TIGIT interactors. Gene Ontology (GO) analysis was used to identify pathways specifically enriched in the CD155-ligated TIGIT interactome (Fig. 2g), identifying two broad categories: signal transduction proteins (including Grb2 and cytoskeletal adapter proteins SdcBP and CD2AP) and endocytosis-related proteins (including TSG101, SNX3 and IST1). To validate candidate proteins, we imaged both IST1 and SdcBP in Jurkat-Raji conjugates (Fig. 2h). Both IST1 and SdcBP were distributed diffusely within the cytoplasm of non-ligated cells but were enriched at the synapse upon CD155-TIGIT ligation in SEE-pulsed Raji cells (Fig. 2h-j). Together, these data provide insights into TIGIT signalling, by highlighting the specific engagement of Ras signalling proteins and endosomal proteins upon CD155 ligation.

**Figure 2.**
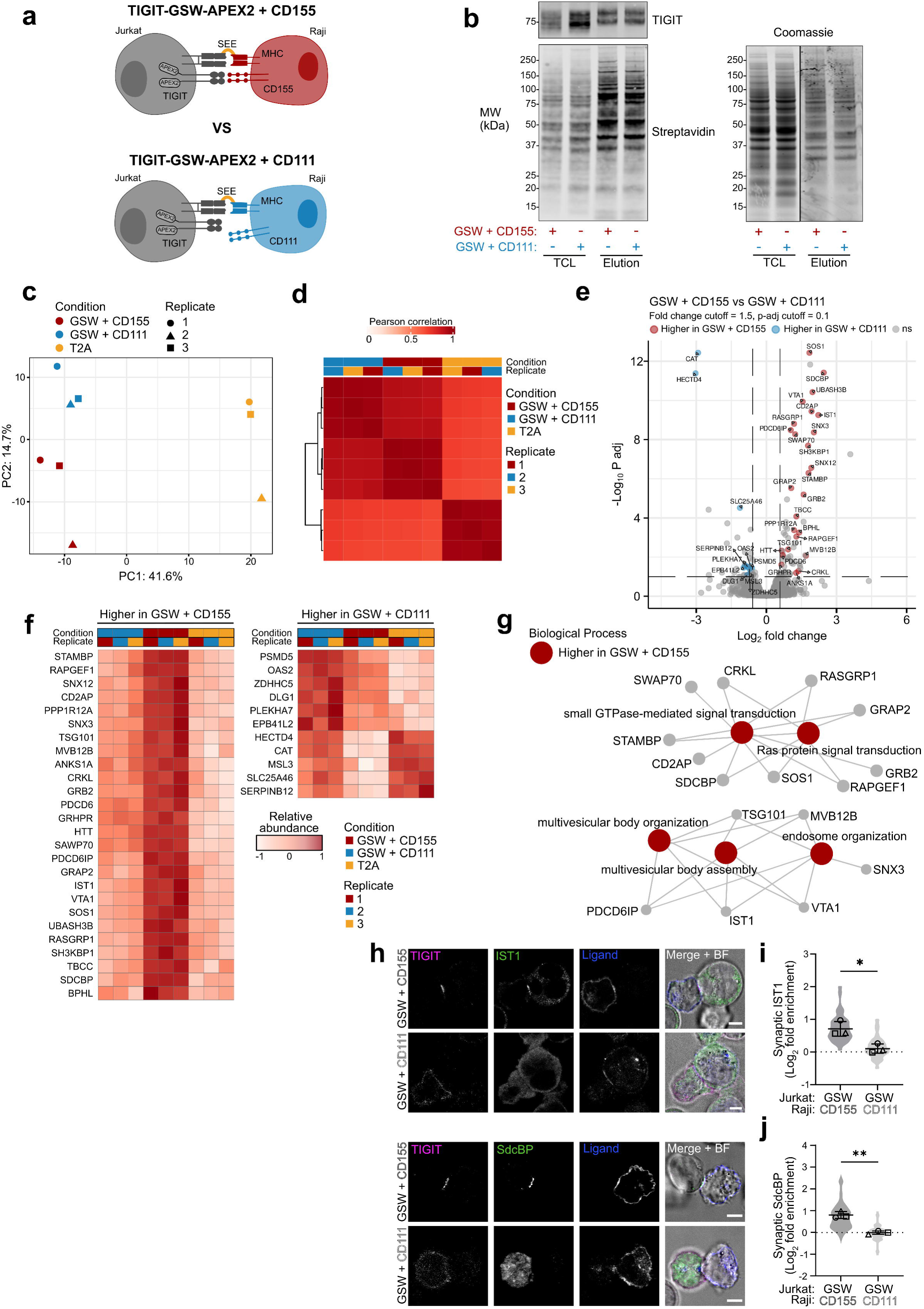
Proximity proteomics reveals the TIGIT interactome following CD155-ligation. **a.** Schematic depicting the cell conjugates used to assess the TIGIT interactome upon ligation with its ligand CD155. TIGIT-GSW-APEX2 (GSW) cells are conjugated with either Raji-CD111 or Raji-CD155 cells in cells stimulated with superantigen (SEE). **b.** Western blot showing TIGIT and biotinylated proteins in GSW Jurkat cells interacting with either Raji-CD111 or Raji-CD155 for 5 mins. We show the banding patterns for both the total cell lysates (TCL) and eluate from streptavidin enrichments, following proximity labelling. Total protein loading controls are shown on the right. Molecular weight markers are indicated on the left. **c.** Principal component analysis depicting the relative positions of each replicate of the conditions analysed according to the variation in the data, with principal components 1 (PC1) and 2 (PC2) plotted. The conditions are colour coded, and replicates are shape coded, as indicated. **d.** Heatmap depicting the overall Pearson correlations between each of the conditions and individual replicates shown in **c**. **e.** Volcano plot showing the differentially proximal proteins to TIGIT in Raji-CD155 conjugates vs Raji-CD111 conjugates. Proteins that were significantly more abundant in Raji-CD155 conjugates are labelled in red and those significantly more abundant in Raji-CD111 conjugates are in blue. Detected proteins not significantly different between the two proteomes are in grey (ns; Cut-off = log_2_ fold change of 1.5, P value = 0.1, as indicated by dashed lines). Both GSW proteomes were pre-filtered against the TIGIT-T2A-APEX2 (T2A) proteome to remove non-specific interactors (Cut-off = log_2_ fold change of 2; P value = 0.05). **f.** Heatmap depicting the measured abundances of the regulated proteins depicted in **e**, abundances are scaled across individual proteins. **g.** GO analysis depicting significantly enriched Biological Process GO terms with the indicated associated proteins within proteins significantly proximal to TIGIT in Raji-CD155 vs Raji-CD111 conjugates. **h.** Confocal microscopy images showing TIGIT (magenta) and either IST1 or SdcBP (green) in GSW expressing Jurkat cells that were conjugated with different Raji cells (expressing either CD111 or CD155) for 5 mins, as indicated on the left of the panel. A V5 stain labels expressed ligands on Raji cells (blue). A merged fluorescence-BF image is provided. Scale bars = 5 μm. **i,j.** Mean log_2_ fold-change in synaptic IST1 (**i**) and SdcBP (**j**) enrichment in Jurkat cells, from the conjugates shown in **h** (±S.D.; n=3 independent experiments; adjusted *P* values from a one-way ANOVA with Tukey’s multiple comparisons are given; violin plots display individual cell variability).

### Ligation-induced interactions require functional inhibitory motifs

As TIGIT significantly relocalises upon ligation (Fig. 1g), we compared the TIGIT interactomes of WT and signalling-deficient forms to delineate which proteins interacted with TIGIT in a signalling-dependent manner from those that were proximal due to localisation changes. To this end, two tyrosine-to-arginine mutations were introduced at positions 225 and 231 in the ITT-like and ITIM inhibitory motifs of TIGIT-GSW-APEX2 (WT), creating YAYA-GSW-APEX2 (YAYA; Fig 3a). Y225 and Y231 are phosphorylated upon ligation and are essential for inhibitory signalling^3,7,11–13^. The mutations did not alter the expression or localisation of TIGIT, as determined by flow cytometry (Fig. 3b) and confocal microscopy (Supplementary Fig. 2a). The YAYA mutant could not be phosphorylated (Supplementary Fig. 2b) and failed to inhibit the release of IL-2 in SEE-pulsed Raji-CD155 conjugates (Fig. 3c).

**Figure 3.**
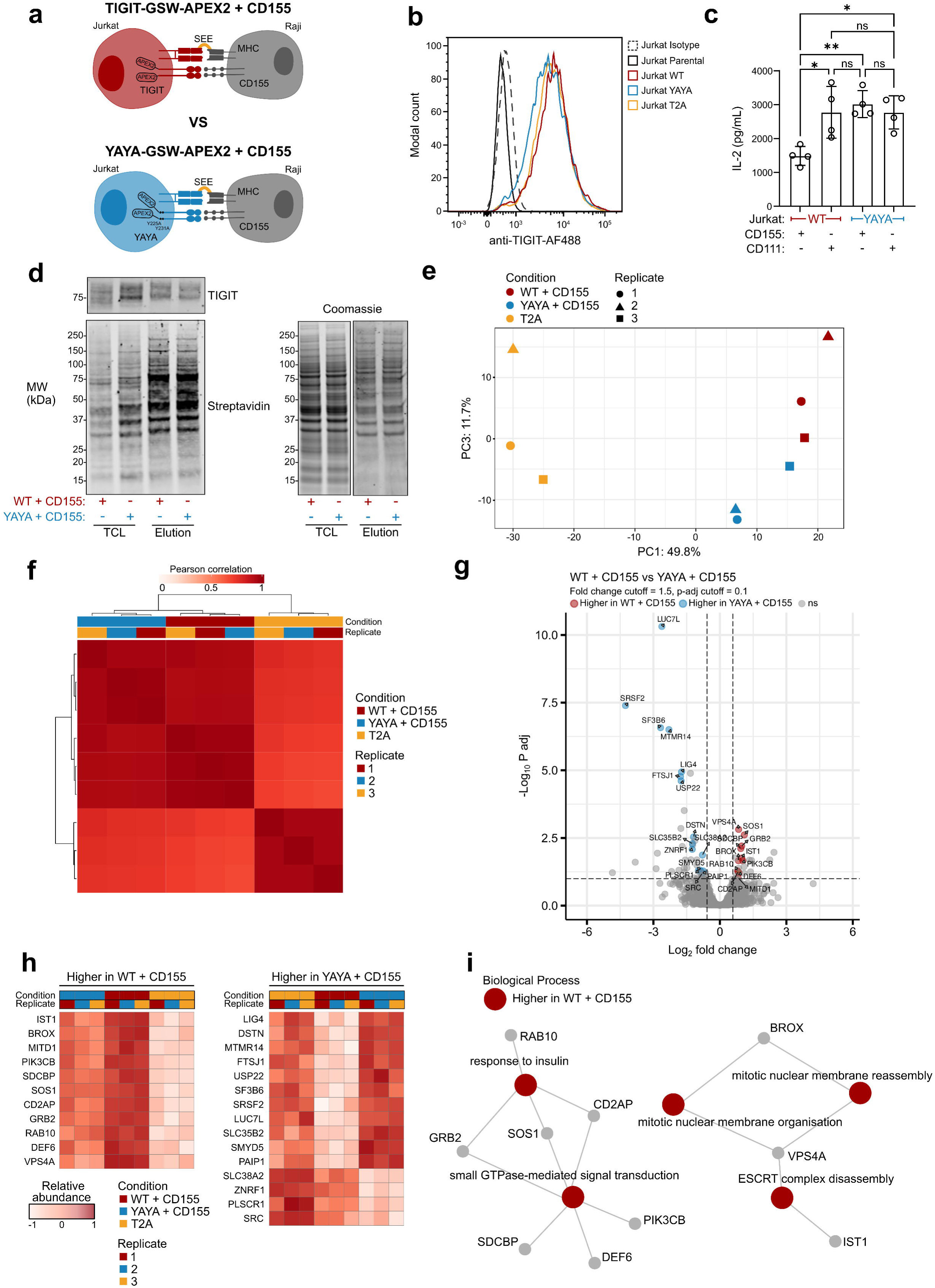
Proximity proteomics reveals the inhibitory motif-dependent TIGIT interactome. **a.** Schematic depicting the cell conjugates used to assess the inhibitory motif-dependent TIGIT interactome. Jurkat cells expressing either TIGIT-GSW-APEX2 (WT) or a signalling-deficient form with Y225A and Y231A mutations, YAYA-GSW-APEX2 (YAYA), were conjugated to CD155-expressing Raji cells pulsed with SEE. **b.** Flow cytometric analysis showing the expression of TIGIT in WT, YAYA, TIGIT-T2A-APEX2 (T2A) and parental Jurkat T cells, with an isotype-matched control. **c.** ELISA data showing the amount of IL-2 released from either WT or YAYA TIGIT-expressing Jurkat cells after co-incubation with SEE-pulsed Raji cells for 6 hrs. (±S.D.; n=4 independent experiments; adjusted *P* values from a one-way ANOVA with Tukey’s multiple comparisons are given; ns = not significant). **d.** Representative western blot showing TIGIT and biotinylated proteins in WT and YAYA TIGIT-expressing Jurkat cells conjugated for 5 mins with SEE-pulsed Raji cells expressing CD155 in both total cell lysates (TCL) and eluate from streptavidin purification, following proximity labelling. A total protein control (Coomassie) is shown to the right from the same TCL and eluates. **e.** Principal component analysis depicting the relative positions of each replicate of the conditions analysed according to the variation in the data, with principal components 1 (PC1) and 3 (PC3) plotted. **f.** Heatmap depicting the overall Pearson correlations between each of the conditions and individual replicates shown in **e**. **g.** Volcano plot showing the differentially proximal proteins to WT TIGIT vs YAYA TIGIT, upon conjugation with SEE-pulsed Raji CD155 cells for 5 mins (Cut-off = log_2_ fold change of 1.5, P value = 0.05, as indicated by dashed lines; colours represent significance). The ligated WT and YAYA proteomes were pre-filtered against the T2A proteome (Cut-off = log_2_ fold change of 2; P value = 0.05). **h.** Heatmap depicting the measured relative abundances of the regulated proteins depicted in **g**, abundances are scaled across individual proteins. **i.** GO analysis depicting significantly enriched Biological Process GO terms with the indicated associated proteins within proteins significantly proximal to WT vs YAYA TIGIT.

Proximity labelling was performed as before, conjugating WT and YAYA Jurkat cells with SEE-pulsed Raji CD155 cells for 5 mins, before inducing labelling (Fig. 3d and Supplementary Fig. 3a, b). Mass spectrometry analysis identified and quantified approximately 2,500 proteins per conjugate and control replicate (Supplementary Fig. 3c). PCA revealed a distinct separation between the T2A control and both the WT and YAYA proximity proteomes in PC1, which explained almost half the variation in the dataset (Fig. 3e). The second and third principal components explained a similar level of variance within the dataset (PC2 = 16.4% and PC3 = 11.7%), with PC2 revealing technical variation and PC3 distinctly separating WT and YAYA proximity proteomes (Fig. 3e and Supplementary Fig. 3d). Pairwise Pearson correlation comparisons indicated both a high similarity within replicate conditions and between the WT and YAYA proximity proteomes (Fig. 3f). Comparing the relative abundances of proteins between the WT and the YAYA mutant proximity proteome identified 11 proteins more proximal to WT TIGIT and 15 proteins more proximal to YAYA (Fig. 3g, h). Of the 11 WT-interacting proteins, almost half were previously enriched in CD155-ligation specific proximity proteomes (Fig. 2f; Grb2, SdcBP, IST1, CD2AP, and Sos1). Similarly, GO analysis of the biological processes enriched within the WT-interacting proteins identified processes associated with the CD155-ligation proximity proteome, including small GTPase-mediated signal transduction components and endocytosis-related proteins (Fig. 3i). Together these data demonstrate that TIGIT selectively engages specific signalling and endocytosis components upon phosphorylation of its inhibitory motifs.

### Inhibitory TIGIT signalling interactions require TCR stimulation

Ligated synaptic TIGIT clusters are nanometres from ligated TCR clusters^11^, and we observed several proteins in the CD155-ligated proximity proteome that are well characterised TCR signalling proteins. Thus, we sought to differentiate TIGIT-specific signalling interactions from those associated with the TCR, ensuring that proteins identified in the TIGIT interactome are not present merely due to their proximity to TCR clusters. We reasoned this could be tested by studying the proximity proteome of GSW cells conjugated with Raji-CD155 cells with or without SEE (Fig. 4a). As before, conjugates were subjected to proximity labelling (Fig. 4b and Supplementary Fig. a, b) and the affinity-purified eluates from conjugates, in addition to the T2A background control, were analysed by mass spectrometry (Supplementary Fig. 4c).

**Figure 4.**
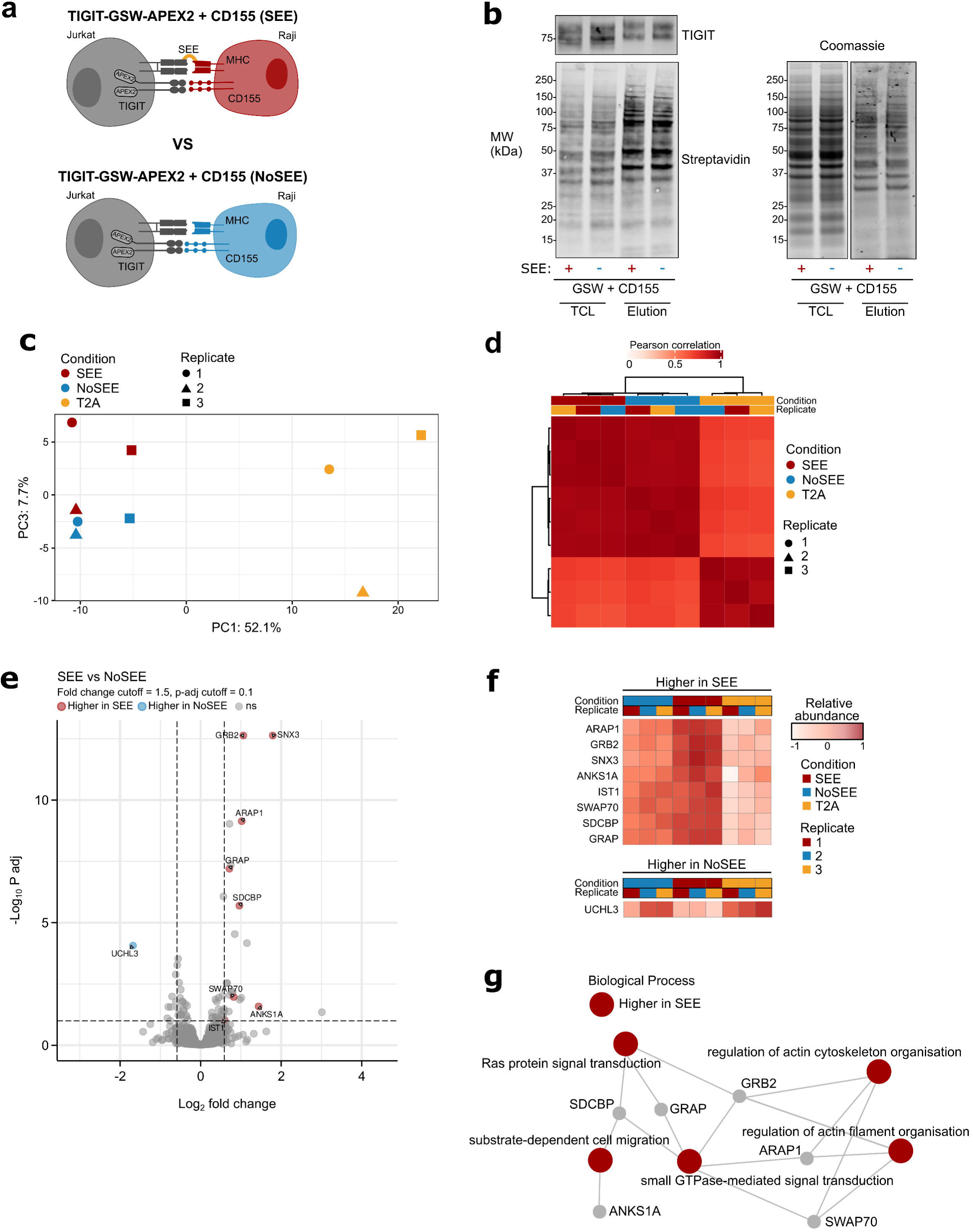
Proximity proteomics reveals that the signalling-specific TIGIT interactome is dependent on T cell activation. **a.** Schematic depicting the cell conjugates used to identify proteins proximal to ligated TIGIT upon T cell activation. Jurkat cells expressing TIGIT-GSW-APEX2 (GSW) were conjugated with Raji CD155 cells with or without SEE-pulsing. **b.** Representative western blot showing TIGIT and biotinylated proteins in GSW Jurkat cells conjugated for 5 mins with Raji CD155 cells with or without SEE-pulsing, in both total cell lysates (TCL) and eluate from streptavidin purification, following proximity labelling. A total protein control (Coomassie) is shown to the right from the same TCL and eluates. **c.** Principal component analysis depicting the relative positions of each replicate of the conditions analysed according to the variation in the data, with principal components 1 (PC1) and 3 (PC3) plotted. **d.** Heatmap depicting the overall Pearson correlations between each of the conditions and individual replicates shown in **c**. **e.** Volcano plot showing the differentially proximal proteins to TIGIT following a 5 mins conjugation with Raji CD155 cells with SEE-pulsing vs without SEE-pulsing (Cut-off = log_2_ fold change of 1.5, P value = 0.05, as indicated by dashed lines; colours represent significance). The ligated TIGIT proteomes were pre-filtered against the T2A proteome (Cut-off = log_2_ fold change of 2; P value = 0.05). **f.** Heatmap depicting the measured relative abundances of the regulated proteins depicted in **e**, abundances are scaled across individual proteins. **g.** GO analysis depicting significantly enriched Biological Process GO terms with the indicated associated proteins within proteins significantly proximal to TIGIT with SEE-pulsing vs without SEE-pulsing.

Again, PCA showed a clear distinction between the replicates of proximity-based interactomes from conjugated GSW cells and the control T2A cells along PC1. The second and third principal components explained 18.2% and 7.7% of the variance in the dataset, respectively, with PC2 separating technical variation and PC3 separating SEE-pulsed and non-pulsed proximity proteomes (Supplementary Fig. 4d and Fig. 4c). Pairwise Pearson correlation comparisons demonstrated significant similarity among replicate conditions, and a high Pearson correlation score observed between the two ligation conditions (Fig. 2d). Comparing the relative abundances of proteins between the SEE-pulsed and non-SEE-pulsed GSW cells revealed 8 proteins more proximal to TIGIT in the SEE-pulsed proximity proteome and 1 protein more proximal to TIGIT in the non-SEE pulsed proximity proteome (Fig. 4e, f). Seven out of the 8 TIGIT interactors more proximal upon T cell stimulation were also more proximal to TIGIT upon CD155-ligation (SNX3, ANKS1A, Grb2, ARAP1, SdcBP, SWAP70, and IST1), and map to similar GO processes (Fig. 4g). Notably, Grb2, SdcBP, and IST1 were identified as proximal to TIGIT in a ligand-, phosphorylation-, and TCR activation-dependent manner. Overall, these findings reveal that TIGIT signalling interactions are dependent on TCR activation, suggesting TIGIT inhibitory function is regulated beyond ligation alone.

### Phosphorylated TIGIT co-localises with Grb2 and CD2AP, but not SdcBP, upon *in vitro* ligation and TCR stimulation

We next sought to validate the increased relative synaptic localisation of TIGIT with Grb2, CD2AP and SdcBP, identified by proximity proteomics, at a higher resolution. PLBs that minimally mimic the different activation states used in the cell conjugation models were generated, using either CD111 or CD155 for ligation and a stimulatory TCR antibody and ICAM-1 for activation. To facilitate the visualisation of Grb2, CD2AP, and SdcBP in Jurkat cells, myc-tagged fusions were introduced into Jurkat TIGIT-SNAP cells (Supplementary Fig. 5). Transduced cells were incubated with PLBs containing ICAM-1 and either CD111 or CD155, or with both CD155 and OKT3 (Fig. 5a). As observed previously, synaptic TIGIT remained diffuse upon contact with PLBs containing CD111, but clustered in the presence of CD155. Grb2 and CD2AP showed diffuse synaptic localisation upon contact with either CD111 plus OKT3 or CD155 without OKT3, but clustered proximal to TIGIT in the presence of both CD155 and OKT3 (Fig. 5a). SdcBP, on the other hand, did not display any noticeable clustering in any condition. To test the phosphorylation-dependency of this interaction, myc-tagged forms of Grb2, CD2AP, and SdcBP were also transduced into Jurkat YAYA-SNAP cells. Limited colocalisation of each myc-tagged protein and YAYA-SNAP was observed in all conditions (Fig. 5b). Quantitative Pearson correlations demonstrated higher correlation coefficients (PCC) between either Grb2 or CD2AP with WT TIGIT when both CD155 and OKT3 were present, but not with YAYA TIGIT or in any SdcBP comparison (Fig. 5c-e).

**Figure 5.**
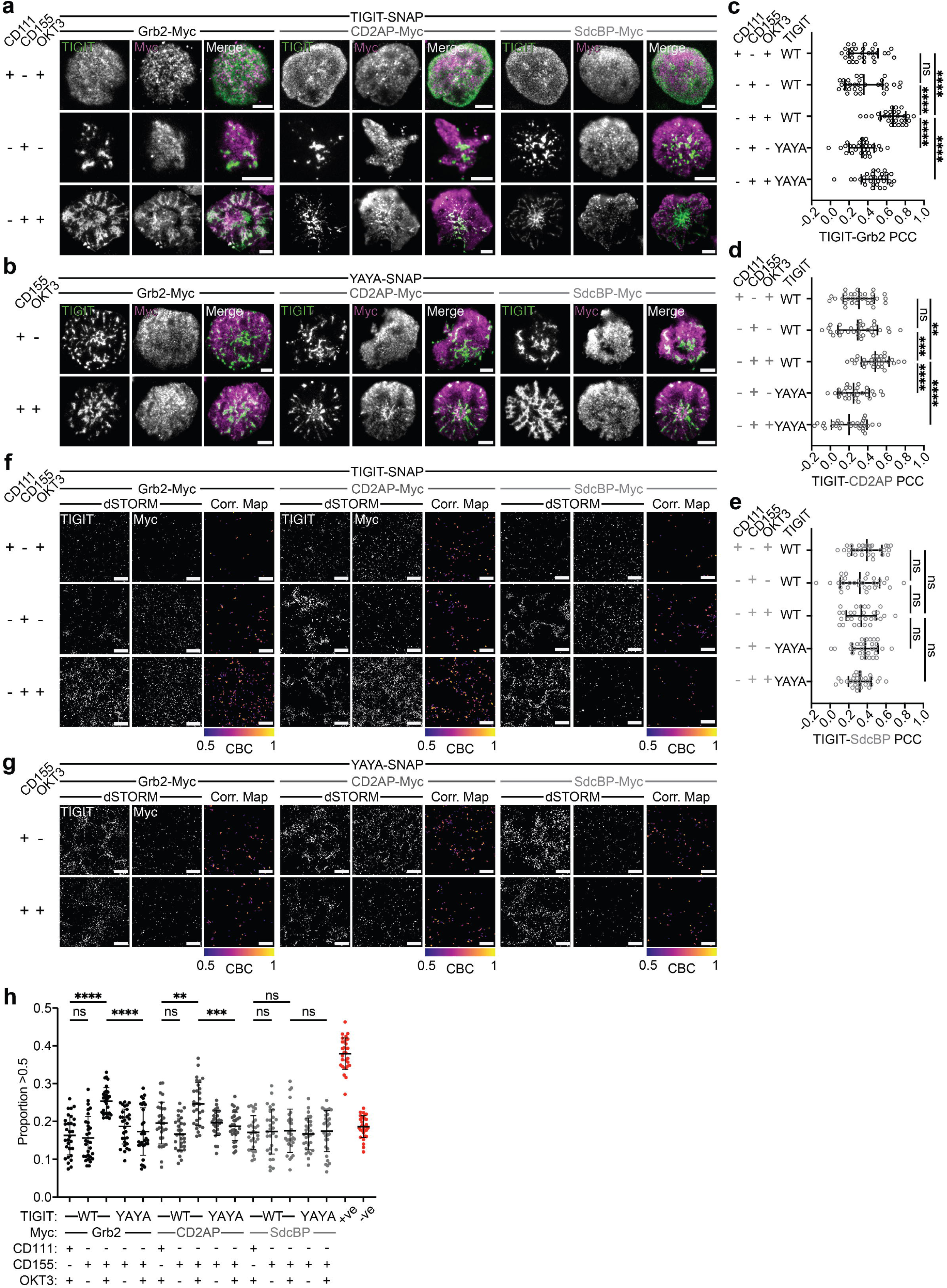
Grb2 and CD2AP, but not SDCBP display nano-proximity to phosphorylated TIGIT clusters following CD155 ligation and TCR activation. a,b. Representative Total Internal Reflection Fluorescence (TIRF) microscopy images of WT (**a**) and YAYA **(b)** TIGIT (magenta) at the immune synapse of TIGIT-SNAP expressing Jurkat cells also expressing one of the indicated myc-tagged proteins (green), that have interacted with planar lipid bilayers (PLB) loaded with ICAM-1 (100 molecules/μm^2^), CD155 or CD111 (400 molecules/μm^2^), and OKT3 (100 molecules/μm^2^), as indicated to the left, for 5 mins. A merged fluorescence image is also provided for each protein comparison, scale bars = 5 μm. **c-e.** Mean Pearson correlation coefficient from a co-localisation analysis of TIGIT with either Grb2 (**c**), SdcBP (**d**) or CD2AP (**e**) from the images shown in **a** and **b** (±S.D.; n = ≥30 individual cells with adjusted P values from a one-way ANOVA with Tukey’s multiple comparisons shown; ns = not significant). **f,g.** TIRF dSTORM images of WT (**f**) or YAYA (**g**) TIGIT and the indicated myc-tagged proteins from the cells displayed in **a** and **b**. Each image represents a smaller 5 μm^2^ region of the indicated TIRF image. For each condition, we also provide a map of TIGIT localisations where coordinate based colocalisation (CBC) values are ≥0.5, and colour code the localisations based on the exact CBC values (Correlation Maps). **h.** Mean proportion of TIGIT localisations with a CBC value ≥0.5 (±S.D.; n = 30 individual cells with adjusted P values from a one-way ANOVA with Tukey’s multiple comparisons shown; ns = not significant).

TIGIT and each myc-tagged protein was then imaged using 2-colour dSTORM to super-resolve the colocalisation we observed upon TCR stimulation and CD155-ligation (Fig. 5f, g). Similar results were observed, with more Grb2 and CD2AP, but not SdcBP, being clustered at the synapse, proximal to TIGIT clusters, upon co-engagement of TIGIT and the TCR (Fig. 5f), except when TIGIT cannot signal (Fig. 5g). As dSTORM is a single molecule localisation technique, it allows a quantification of the inter-molecular distances. Thus, we used coordinate based colocalisation (CBC^18^) and nearest-neighbour distance to quantify the extent of colocalisation, using a primary vs. secondary antibody staining of TIGIT as a positive control, and inverting the XY coordinates of the secondary stain to generate a comparable negative control (Supplementary Fig. 6). By plotting the TIGIT localisations with CBC values ≥0.5, the extent of colocalisation between TIGIT and each myc-tagged fusion protein can be observed, demonstrating higher colocalisation when WT TIGIT and the TCR are co-engaged for Grb2 and CD2AP (Fig. 2f, g), that were significantly different from all other conditions tested (Fig. 2h). The lack of activation induced TIGIT and SdcBP colocalisation was surprising as SdcBP was observed to colocalise with TIGIT at the IS upon CD155 ligation and superantigen stimulation in a cell conjugate model (Fig. 2h). This suggests additional interactions between Raji and Jurkat cells, not present in our minimal *in vitro* model, could drive SdcBP recruitment, or that the introduction of the Myc tag to SdcBP disrupted SdcBP-TIGIT interactions. Overall, ligation of TIGIT and the TCR using an *in vitro* artificial APC promotes nanoscopic TIGIT-Grb2 and TIGIT-CD2AP interactions in an inhibitory signalling-dependent manner, validating CD2AP as a novel TIGIT interactor.

### TIGIT internalisation is phosphorylation- and TCR activity-dependent

Given that we observed increased proximity of TIGIT to endocytosis-related proteins upon ligation, T cell activation and phosphorylation, we aimed to test whether TIGIT internalisation was also dependent on these factors. To test this, changes in total and surface TIGIT abundance were measured in TIGIT-SNAP Jurkat cells expressing either WT (WT-SNAP) or signalling-deficient TIGIT (YAYA-SNAP), upon conjugation with Raji-CD155 cells. Flow cytometry quantified the surface TIGIT abundance by antibody staining, and the total TIGIT abundance through SNAP-tag labelling prior to conjugation, in different conjugation conditions (Fig. 6a and Supplementary Fig. 7a-d). Conjugation to Raji-CD155 cells, in the presence or absence of superantigen stimulation, did not induce changes to the total TIGIT abundance (Fig. 6a, b). However, SEE-stimulation in CD155-ligated cells induced a significant decrease in surface TIGIT (Fig. 6a, c), which was not observed with the signalling-deficient form. This decrease was significantly reduced at 4°C (a common method to slow internalisation) and was reduced by Src-family kinase inhibitors Dasatinib and PP2 (Fig. 6a-c). As expected, surface TIGIT remained unchanged when ligated with Raji-CD111 cells (Supplementary Fig. 7e, f). Together, this shows that TIGIT internalisation is dependent on its phosphorylation and requires both ligation and TCR activation.

**Figure 6.**
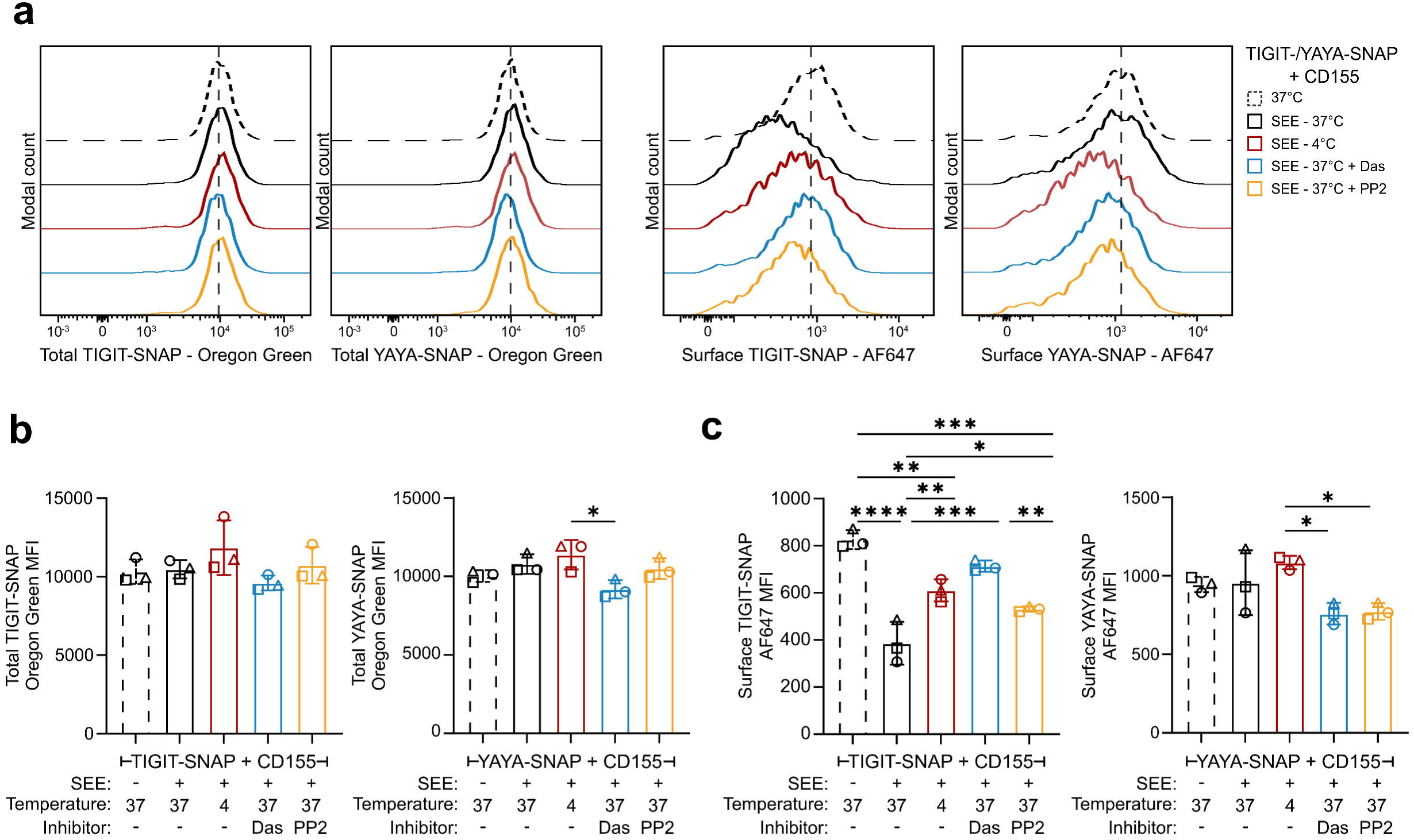
TIGIT internalisation is dependent on both CD155-ligation, TCR activation and TIGIT phosphorylation. **a.** Histograms showing the levels of total TIGIT (left two graphs; labelled with Oregon Green SNAP ligand) and surface TIGIT (right two graphs; labelled with an AF647-conjugated TIGIT antibody) in TIGIT-SNAP-(WT) or YAYA-SNAP mutant-expressing Jurkat cells, as indicated. Conditions are indicated to the right. **b, c.** Mean total (**b**) or surface (**c**) TIGIT abundance in TIGIT-SNAP (left) or YAYA-SNAP (right) Jurkat cells in each of the indicated conditions, as shown in **a** (±S.D.; n = 3 independent experiments with adjusted P values from a one-way ANOVA with Tukey’s multiple comparisons shown).

### TIGIT phosphorylation is dependent on CD155 ligation and TCR activation

Proximity proteomics and super-resolution imaging demonstrated that TIGIT signalling associations were dependent on both CD155 ligation and TCR stimulation. Furthermore, all associations discovered thus far were also dependent on TIGIT phosphorylation. While we expected signalling proteins to interact in a CD155 ligation- and phosphorylation-dependent manner, the requirement for TCR activation was surprising. This led us to question whether TIGIT phosphorylation itself also required TCR activation. To assess this, Jurkat TIGIT-SNAP cells were conjugated to Raji-CD111 or CD155 cells for 5 mins in the presence or absence of superantigen. TCR activation occurred in both conjugates where SEE was present, as assessed by phosphorylated CD3ζ (Fig. 7a). We assayed TIGIT phosphorylation in each condition using Phos-tag SDS-PAGE and observed a mobility shift, representing the phosphorylated form of TIGIT, only following conjugation with SEE-pulsed Raji-CD155 cells (Fig. 7a, b). Thus, although ligation with CD155 can cause TIGIT to enrich and cluster at immune synapses, this does not induce its phosphorylation. This could feasibly be accounted for by an increase in cell conjugations in the presence of superantigen. To test this hypothesis, Jurkat and Raji cells were labelled, conjugated for 5 mins, fixed and analysed by flow cytometry to determine the number of conjugates (Fig 7c). Indeed, CD155-ligation approximately doubled cell conjugations, but the presence of superantigen had no effect. Thus, TIGIT phosphorylation was not a result of more cellular interactions but likely mediated by increased local kinase activity generated by TCR stimulation. Previous work *in vitro* has demonstrated TIGIT phosphorylation by Src-family kinases^7,13^, known to be active at activated T cell synapses^19,20^. We therefore investigated whether TIGIT phosphorylation was impacted by Src-family kinase inhibitors and found that both Dasatinib and PP2 could prevent TIGIT phosphorylation (Fig. 7d). However, decreases in TIGIT phosphorylation induced by these kinase inhibitors were proportional to their ability to reduce TCR phosphorylation (Fig 7d), and thus more work will be required to elucidate whether Src-family kinases directly mediate TIGIT phosphorylation. Overall, we show that TIGIT phosphorylation is dependent on both CD155-ligation and TCR activation and can be inhibited by Src-family kinase inhibition.

**Figure 7.**
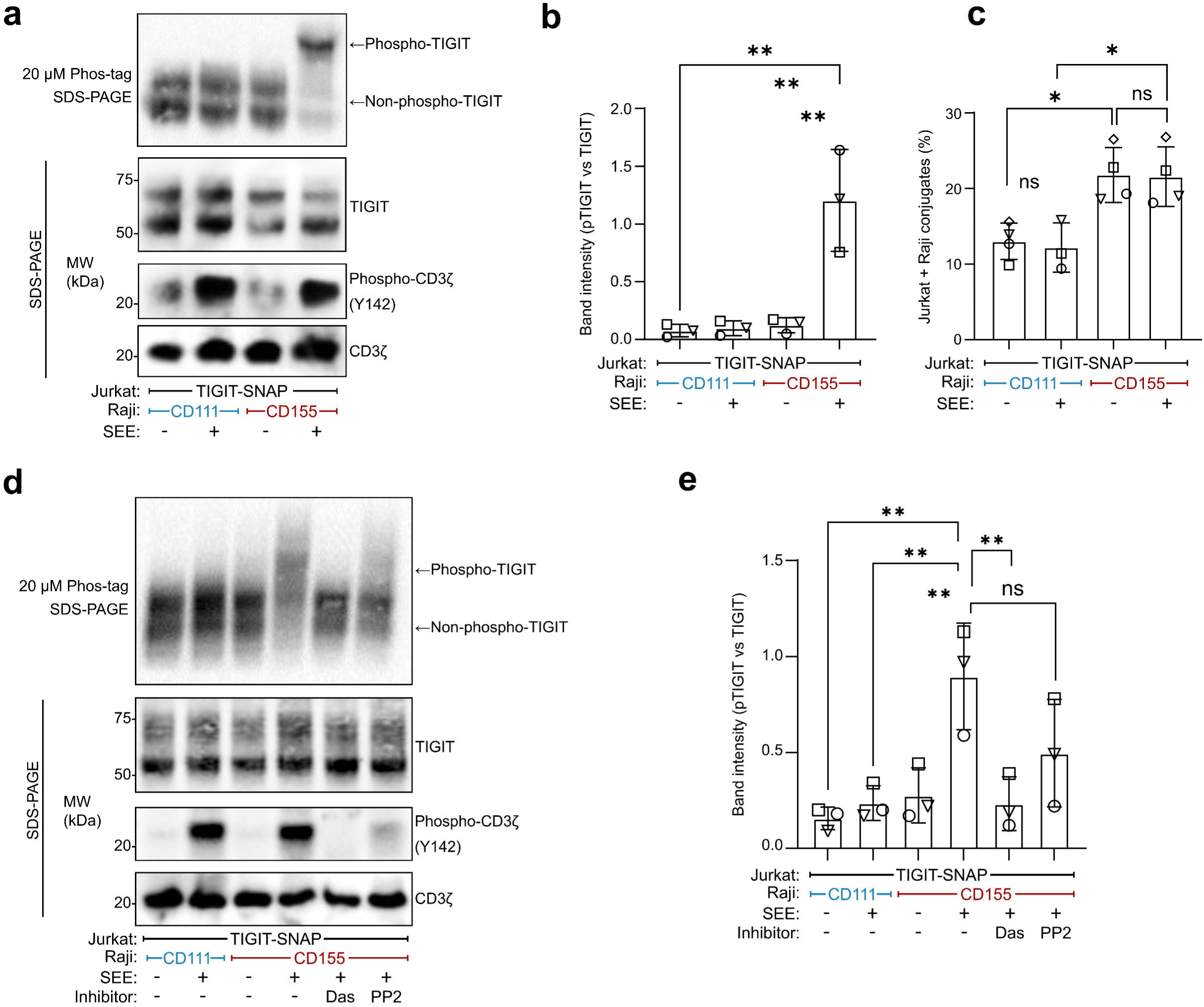
TIGIT phosphorylation is dependent on both CD155-ligation and TCR activation and is mediated by a Src-family kinase. **a.** Representative western blot analysis of TIGIT using either Phos-tag SDS-PAGE (top) or standard SDS-PAGE (middle) to examine TIGIT phosphorylation in Jurkat TIGIT-SNAP cells conjugated to Raji CD155 or CD111 cells for 5 mins, as labelled below. SEE addition to stimulate the TCR is also indicated below. The abundance of both phosphorylated CD3ζ (bottom; standard SDS-PAGE) and total CD3ζ was assessed to evaluate T cell activation. **b.** Quantification of band intensity ratio between phosphorylated and non-phosphorylated TIGIT from Phos-tag SDS-PAGE images shown in **a** (±S.D.; n = 3 independent experiments with adjusted P values from a one-way ANOVA with Tukey’s multiple comparisons shown). **c.** Flow cytometric analysis showing the number of conjugated Jurkat TIGIT-SNAP cells (stained with CFSE) and Raji CD155 or Raji CD111 cells (stained with CTV), with or without SEE (±S.D.; n = 4 independent experiments with adjusted P values from a one-way ANOVA with Tukey’s multiple comparisons shown; ns = not significant). **d.** Representative western blot analysis of TIGIT using either Phos-tag SDS-PAGE (top) or standard SDS-PAGE (bottom) to examine TIGIT phosphorylation in Jurkat TIGIT-SNAP cells conjugated to Raji CD155 or CD111 cells, in the presence of the indicated kinase inhibitors (Das; Dasatinib and PP2) and/or SEE. **e.** Quantification of band intensity ratio between phosphorylated and non-phosphorylated TIGIT from Phos-tag SDS-PAGE images shown in **d** (±S.D.; n = 3 independent experiments with adjusted P values from a one-way ANOVA with Tukey’s multiple comparisons shown).

## Discussion

Here, we show that TIGIT engages various signalling molecules upon ligation, including the previously identified interactor Grb2 and novel interactors such as the cytoskeletal adaptor proteins SdcBP and CD2AP. TIGIT also engaged proteins from endosomal pathways including IST1 and SNX3. These interactions were driven by phosphorylation of tyrosines Y225 and Y231 in its inhibitory motifs, but unexpectedly also required concurrent TCR stimulation. Textbook depictions of receptor signal transduction, including those of immune checkpoint receptors, present linear cascades where ligation drive conformational changes and clustering to mediate activation and signalling. Yet here, TIGIT phosphorylation, which controlled its internalisation and signalling, only occurred with both CD155 ligation and TCR stimulation. In other words, TIGIT ligation by itself was unable to elicit TIGIT signalling, despite this leading to extensive clustering at the immune synapse. When both TIGIT and the TCR are engaged, they coalesce within nanometre-scaled, dense clusters. TCR clustering and activation generates a Src-family kinase-rich, phosphatase-poor local environment^20^, which we propose provides the favourable biochemical conditions for TIGIT phosphorylation, and subsequent signalling.

The requirement for both ligation and TCR activation to drive inhibitory receptor signalling is likely not unique to TIGIT. PD-1 also requires proximity to TCR clusters for phosphorylation and function^21^. Chimeric PD-1 molecules containing bulky extracellular domains can cluster upon PD-L1 ligation but are excluded from ligated TCR nanoclusters. Exclusion resulted in reduced PD-1 phosphorylation, recruitment of the phosphatase Shp2 and subsequent inhibition of IL-2 secretion. The shared dependence of PD-1 and TIGIT on proximal TCR activation may explain their effectiveness as combination therapy targets, as both are active in the same spatially defined regions of the cell and may inhibit similar processes. The contextual activation of an immune checkpoint, such as TIGIT, could ensure inhibition is only triggered when required^16^. This could reduce the risk of unwarranted immune suppression and receptor internalisation in the absence of activation, thus retaining function when required.

The co-proximity of checkpoints to stimulatory receptors is also an important consideration for therapies. A recent study enhanced PD-1 blockade by forcing PD-1 to interact *in cis* with the bulky extracellular phosphatase CD45 through a bi-specific molecule^22^. This process, termed Receptor Inhibition by Phosphatase Recruitment (RIPR), excluded PD-1 from the immune synapse and its kinase-rich environment. This spatial isolation prevented PD-1 from interacting with and regulating CD28, through Shp2 recruitment^23^, while also forcing PD-1 into the proximity of a promiscuous phosphatase reducing tonic phosphorylation. Forcing PD-1 and/or TIGIT to interact *in cis* with a bulky receptor that isn’t a phosphatase, such as CD43^24^, may also induce a similar phenomenon, given that receptor phosphorylation is linked to its sub-synaptic localisation. This approach opens new avenues for blockade, and, simultaneously, offers strategies to drive potent agonism by forcing immune checkpoint receptors to localise to stimulatory clusters.

This study also sheds light on the potential mechanisms driving TIGIT inhibition. By systematically employing proximity proteomics in different conjugation conditions, we were able to map TIGIT interactomes resulting from ligation, and determine which interactions were specific to its inhibitory signalling motifs within the context of T cell activation. In NK cells, the adaptor protein Grb2 can interact with phosphorylated Y225 and Y231 in the ITT-like and ITIM motifs^7,13^. This interaction has proved difficult to capture in T cells and has only been observed with peptide purifications in cell-free lysates^3^. Proximity proteomics can more readily identify labile or transient interactions, compared to precipitation-based approaches^25^. Here, it was able to detect the TIGIT-Grb2 interaction *in situ*, providing validation that Grb2 is a bone fide

TIGIT interactor in T cells. Grb2 can recruit SHIP-1, blunting downstream MAPK, Akt and NF-κB signalling^13,14^. However, Jurkat cells do not express SHIP-1^26^, demonstrating that TIGIT can signal to inhibit independently of this phosphatase. GO analysis of TIGIT interactors consistently identified cytoskeletal and vesicular processes, through molecules such as SdcBP, CD2AP and IST1. Cytoskeletal processes are critical in mediating immune synapse formation, synaptic signalling and cytolytic function^27–32^ and inhibitory receptors have been implicated to disrupt their synaptic function^33,34^. Similarly, vesicular processes are also important in mediating synaptic function and can regulate TCR activation^35–38^. Thus, TIGIT may function by disrupting the cytoskeletal or trafficking processes that mediate signalling at the synapse. We show TIGIT phosphorylation is required for its internalisation, which correlates with its ability to inhibit. Internalisation may inhibit T cell activity by: i) reducing CD155 levels on APCs by trans-endocytosis to prevent CD226 signalling, ii) disrupting TCR signalling through the displacement of key signalling molecules (like Grb2) from the synapse or iii) displacement of the TCR itself from the synapse. These novel interactions will require future investigation to elucidate their roles in the inhibitory function of TIGIT.

Collectively, this study advances our understanding of how TIGIT functions in T cells, demonstrating its context-specific activation. TIGIT signalling is not triggered by ligation alone but requires concomitant activation of the TCR. As numerous inhibitory receptors localise to TCR clusters upon co-ligation, this suggests the regulation of inhibitory receptor activity by TCR signalling is broadly important for checkpoint activation.

## Methods

### Cell lines

Jurkat cells were obtained from the ATCC (Clone E6-1; ATCC^®^ TIB-152™; RRID:CVCL0367) and maintained in RPMI-1640 Medium (R0883; Sigma Aldrich) supplemented with 2 mM L-glutamine (25030081; Thermo Fisher Scientific), 10% heat-inactivated fetal bovine serum (FBS; A5256801; Sigma Aldrich), and 50 U/mL Penicillin-Streptomycin (P/S; 15140122; Thermo Fisher Scientific). Jurkat TIGIT-SNAP and YAYA-SNAP cells were generated previously^11^. Raji cells were obtained from the ATCC (ATCC^®^ CCL-86; RRID:CVCL0511) and maintained as per Jurkat cells. HEK293T cells were obtained from the ATCC (HEK 293T/17; ATCC^®^ CRL-11268; RRID:CVCL1926) and maintained in DMEM, High Glucose (41965039; Life Technologies) supplemented with 10% heat-inactivated FBS and 50 U/mL P/S. All cells were maintained at 37°C, 5% CO_2_ within tissue culture incubators.

### Molecular cloning and plasmid generation

pLX304 TIGIT-GSW-APEX2 and pLX304-TIGIT-T2A-APEX2 was generated by sequential cloning. First, an amplified PCR product of the TIGIT gene (from pENTR/Zeo-TIGIT-NoSTOP^11^) was inserted into BstBI digested pLX304-Flag-APEX2-NES (a gift from Alice Ting; Addgene plasmid # 92158; http://n2t.net/addgene:92158; RRID:Addgene_92158^39^) using HIFI DNA assembly (E2621; New England Biolabs) to create pLX304-TIGIT-Flag-APEX2-NES. Next, either the T2A cleavage peptide or the GGSx2 + Waldo Linker (GSW linker) was amplified from previously generated plasmids and inserted into BamHI-digested pLX304-TIGIT-Flag-APEX2-NES constructs to generate either pLX304 TIGIT-T2A-APEX2 or pLX304-TIGIT-GSW-APEX2.

Myc-tagged Grb2, CD2AP, and SdcBP TIGIT-SNAP Jurkat cells were generated in house using a lentiviral vector that includes a N-terminally myc tagged gene with a doxycycline inducible promoter as the backbone. Briefly nucleotide gene strings (GeneArt, Thermo Fisher Scientific), were designed for Grb2, CD2AP, and SdcBP with overhangs that matched the destination vector. Overhang at the start of the string was GGCGGCGGAGGATCTACCGGT and at the end of the string was GAATTCTAAAAGGATCTGCG. The gene strings were combined into the linearised backbone vector with HIFI DNA assembly.

YAYA-GSW-APEX2 mutant was generated using pLX304 TIGIT-GSW-APEX2 as a backbone. Primers were designed to generate two fragments from restriction enzymes, which are linked by primers designed over the amino acid region 222-233 and designed to introduce tyrosine-to-alanine mutation at 225 and 231 to generate the mutations. Primers were GTACATCAAGTGTATCATATGCCAAGTACGCCCCC (TIGIT forward), GCTTCTG<u>CG</u>ACTCAGGACATTGAAG<u>CG</u>GTCATGC (YAYA reverse), GCATGAC<u>GC</u>CTTCAATGTCCTGAGT<u>GC</u>CAGAAGC (YAYA forward), and CGCGCCACCGGTTAGCGCTAGCTCATTACTACGTAGAATC (APEX2 reverse). Primers were amplified and combined into the linearised backbone vector with HIFI DNA assembly.

### Lentiviral production, transduction, and selection of stable cell lines

To generate lentiviral vectors, 18 x 10^6^ HEK293 cells were seeded onto a T175 flask for each transfection condition. 20 h later, 95 μg of total DNA was diluted in Opti-MEM (31985; Thermo Fisher Scientific) at a ratio of lentiviral construct (pLX or pCDH) to viral packaging construct (psPAX2; Addgene plasmid #12260) to viral envelope construct (pMD2.G; Addgene plasmid #12259) in a 4:3:3 (w/w/w). Polyethylenimine MAX 40K (PEI) (24765; Polysciences Catalog) was incubated with 6 mL of Opti-MEM at 3x the concentration of DNA (18 μg) for 2 mins at room temperature (RT). The DNA/Opti-MEM and PEI/Opti-MEM mixture were vortexed together and incubated for 20 mins at RT. After removing the media from the HEK293 cells, the Opti-MEM mixture was carefully dripped onto the cells and incubated at 37°C with 5% CO_2_ for 6-8 h. After incubation, the media was replaced, and the cells were incubated for 48 h. The harvested supernatants were centrifuged and passed through a 0.45 μm cellulose acetate filter (E4780-1453; Starlab) to remove cell debris. The virus was concentrated by ultracentrifugation at 35,000 g for 3 h at 4°C and the pellets were resuspended in 110 μL Opti-MEM. 5-20 μL of the virus in Opti-MEM was transduced into Jurkat cells for one week and selected using 5 μg/mL puromycin.

### Activation of Jurkat cells with Staphylococcal Enterotoxin E (SEE)-pulsed Raji cells

Raji cells were counted and washed in media. SEE (ET404; Toxin Technologies) was added to Raji cells at 60 ng/mL and incubated at 37°C, 5% CO_2_ for 30 mins. Excess SEE was washed with complete media and the cells resuspended in media. Jurkat cells were counted and mixed with Raji cells at a ratio of Jurkat:Raji of 2:1 and centrifugation-assisted conjugation was performed by spinning at 50 g for 30 seconds. The cells were incubated at 37°C, 5% CO_2_ for 6 h before the supernatants were collected following centrifugation at 500 g for 5 mins to remove cells.

### Flow cytometry

To assess the abundance of cell surface proteins, cells were stained with live/dead Fixable Blue (L23105; Thermo Fisher Scientific) for 15 mins at RT. After washing, cells were blocked with 1% of human serum (HS; H4522; Sigma Aldrich) for 10 mins at RT, followed by antibody staining for 30 mins at 4°C. Finally, cells were washed and fixed in 4% paraformaldehyde (PFA; 28908; Thermo Fisher Scientific) for 20 mins at 4°C, before being analysed by flow cytometry (BD FACS LSRFortessa X20).

To assess the number of conjugates in co-cultured cells, Jurkat and Raji cells were either stained with CellTrace CFSE (C34554; Thermo Fisher Scientific) or CellTrace Violet (C34557; Thermo Fisher Scientific) cell proliferation kit for 15 mins at 37°C. The Raji cells were then incubated with and without SEE for 30 mins at 37°C, 5% CO_2_ and washed in complete media. The Jurkat and Raji cells were conjugated by centrifugation at 50 g for 30 seconds and incubated at 37°C, 5% CO_2_ for 5 mins. The cells were fixed in 4% PFA for 20 mins and analysed by flow cytometry (BD FACS LSRFortessa X20).

To measure the internalisation of TIGIT, TIGIT-SNAP and YAYA-SNAP cells were initially incubated with media containing 5 μM SNAP-Cell Oregon Green (S9104S; New England Biolabs) for 30 mins at 37°C, 5% CO_2_. Washed three times with complete media and incubated for another 30 mins at 37°C, 5% CO_2_ to remove unreacted SNAP-tag substrate, before a final wash. Raji cells were pre-pulsed with SEE as described above. Jurkat cells treated with inhibitors were incubated with 250 nM of Dasatinib (SML2589; Sigma Aldrich) or 50 μM of PP2 (P0042; Sigma Aldrich) for 30 mins at 37°C, 5% CO_2_. The Jurkat and Raji cells were conjugated at a ratio of 1:1 by centrifugation at 50 g for 30 seconds and either incubated on ice or 37°C, 5% CO_2_ for 30 min. Following incubation all samples were placed on ice and all centrifugation steps were performed at 4°C. The cells were resuspended in LIVE/DEAD Fixable Near IR (780) Viability Kit (L34994; Thermo Fisher Scientific) for 15 mins at 4°C. Subsequently, the cells were blocked with 3%BSA/1%HS for 15 mins at 4°C before the antibodies were spiked in. The cells were washed 3x with FACs buffer (2% BSA, 5 mM EDTA (E8008; Sigma Aldrich)) and fixed and permeabilised using the eBioscience Fixn/Perm Buffer Set (88-8824-00; Thermo Fisher Scientific). Finally, the cells were washed with 3x FACs buffer and analysed by flow cytometry (BD FACS LSRFortessa). Analysis of flow cytometry data was performed on FlowJo™ v10.10 software.

### Antibodies

The antagonistic TIGIT antibody which blocked TIGIT-CD155 was added to Jurkat cells at 5 μg/mL on ice, 10 mins prior to use in cytokine secretion assays (Clone VSIG9.01; produced by the Center for proteomics, Faculty of Medicine, University of Rijeka). For isotype controls, the clone MOPC-21 (400166; Biolegend) or clone G3A1 (5415; Cell Signaling Technology) was used. For western blot the following antibodies were used: TIGIT (E5y15; 1:2000; Cell Signaling Technology), CD3ζ (6B10.2; 1:1000; sc-1239; Santa Cruz Biotechnology), pCD3ζ (Y142; 1:1000; K25-407.69; 558402; BD Biosciences), α-tubulin (DM1A; 1:1000; 62204; Thermo Fisher Scientific), myc (9E10; 1:1000; Abcam), streptavidin Alexa Fluor 680 (S21378; 1:10000, Thermo Fisher Scientific), αRabbit IgG DyLight™ 800 Conjugate (SA5-10036; 1:10000, Thermo Fisher Scientific), αRabbit IgG-HRP (#7074; Cell Signaling Technology) and αMouse IgG-HRP (7076; Cell Signaling Technology). For immunofluorescence and flow cytometry experiments the following antibodies were used: αTIGIT (MBSA43; 2.5 μg/mL), αV5 (Rabbit polyclonal; NB600-381; Novus Biologicals; 1 μg/mL), αCD19 (HIB19; 740287; BD Biosciences; 1 μg/mL), αCD3ε (SK7, 344822; Biolegend; 1 μg/mL), αIST1 (51002-1-AP, Proteintech, 2.5 μg/mL), αSdcBP (68096-1-Ig; Proteintech, 2.5 μg/mL), NeutrAvidin (22832; Thermo Fisher Scientific; 2.5 μg/mL), and myc (9E10; 626810/ab206486; BioLegend/Abcam, 2.5 μg/mL). OKT3 was monobiotinylated according to the protocol previously published^40^ using EZ-link™ Sulfo-NHS-LC-LC-biotin (21338; Thermo Scientific; Biotin reagent solution used at 0.1 μg/mL with the antibody at 1mg/mL in Dulbecco′s Phosphate Buffered Saline (DPBS; D8537; Sigma Aldrich). Antibodies that were not conjugated directly from the purchaser were then conjugated in house with NHS-Esters of each dye (Alexa Fluor 647; A20006; Invitrogen, Atto488; 41698; Sigma Aldrich) and desalted with Zeba™ spin columns (7K MWCO; 89882; Thermo Scientific).

### Confocal Imaging

Nunc™ Lab-Tek™ II chambered coverglass (155409; Thermo Fisher Scientific) were pre-coated with 0.01% (v/v) poly-L-lysine (P78890; Sigma Aldrich) at RT for 10 mins and then 5 μg/ml fibronectin (F0985; Sigma Aldrich) for 1 h at 37°C, 5% CO_2_. Raji cells were pre-pulsed with SEE following the previously stated method. For standard co-culture imaging, Jurkat and Raji cells were co-cultured at a ratio of Jurkat:Raji of 1:1 by centrifugation at 50 g for 30 seconds and then gently transferred to the Lab-Tek™ for 5 mins. For co-culture proximity labelling imaging, the ligand-expressing Raji cells were added to Jurkat cells that have been incubated with biotin-phenol (BP; SML2135; Sigma Aldrich) for 15 mins, centrifuged for 30 seconds at 50 g and incubated for 2 mins at 37°C, 5% CO_2_. These cells were gently transferred to the Lab-Tek™ plate for 13 mins at 37°C, 5% CO_2_. Subsequently, hydrogen peroxide (H_2_O_2_) (386790; Sigma Aldrich) was added for 2 mins at RT. Immediately after incubation, samples were placed on ice and washed three times with quenching solution (20 mM sodium ascorbate (PHR1279; Sigma Aldrich), 10 mM Troxol (238813; Sigma Aldrich) and 20 mM sodium azide (S2002; Sigma Aldrich) in DPBS). Both proximity-labelled and non-proximity labelled co-cultured cells were subsequently fixed with 4% PFA for 20 mins at 37°C. The wells were permeabilised with 0.1% Triton X-100 (X100; Sigma Aldrich) for 10 mins at RT, then blocked overnight at 4°C with blocking buffer (3%BSA/1%HS). Samples were washed in DPBS and stained with primary antibody in blocking buffer for 1 h at RT. If a secondary antibody was required, samples were washed with DPBS, and the secondary antibody in blocking buffer was added for 1 h at RT. Samples were washed with DPBS and stored in DPBS for imaging.

Confocal microscopy (Leica TCS SP8) was performed using a 100x/1.40 NA oil-immersion objective and white light laser source. Images were analysed using Fiji, which determined receptor accumulation at the synapse and the level of biotinylation. Receptor accumulation was calculated by dividing the fluorescence intensity at the cell-contact interface with the fluorescence intensity across the rest of the cell.

### TIRF and Single Molecule Localisation Microscopy

TIRF microscopy was performed on an inverted microscope (Leica SR 3D-GSD) fitted with a HC PL APO 160X oil immersion lens (NA 1.43), an EMCCD camera (Andor iXon Ultra 897), and the TIRF laser penetration depth set to 150 nm. All TIRF images were recorded using the same acquisition settings (exposure duration and laser power) and imaged in DPBS. Colocalisation analysis by Pearson correlation of two individual channels was carried out in Fiji. Firstly, regions of interest (ROI) were drawn around the cells (aided by the threshold function and inspected manually), and PCCs of the two channels (TIGIT and myc) within the ROIs were calculated using the Coloc-2 plugin. All coefficients presented derive from non-thresholded comparisons.

Dual colour STORM images were acquired sequentially, firstly by excitation with the 647-nm laser (15%), acquiring 7,500 frames with a 11-ms exposure time and followed by excitation with the 488-nm laser (50%), acquiring 7,500 frames with a 11-ms exposure time and an electron multiplier gain set to 120. For both colours, we boosted recovery of fluorophores to the On state through excitation with a 405-nm laser (5%) after the first 1,000 frames. Samples were immersed in 0.22 μm-filtered OxEA imaging buffer (50 mM β-MercaptoEthylamine hydrochloride (MEA; 30078; Sigma Aldrich), 3% (v/v) OxyFluor™ (OF; Oxyrase Inc.), 20% (v/v) sodium DL-lactate solution (L4263; Sigma Aldrich) in DPBS, pH adjusted to 8–8.5 with NaOH), as previously published^41^.

Molecules were localised and images reconstructed using the ThunderSTORM software^42^. Raw images were firstly filtered to remove noise and enhance blinking ‘events’ from single fluorophores (wavelet filter B-spine method, order 3, scale 2). Initial event detection was determined using the local maximum localisation method, with a threshold for the peak intensity set to 2 times the standard deviation of the F1 wavelet, and 8-neighborhood connectivity for each pixel screened. Subpixel event localization was calculated using an integrated Gaussian point-spread function and maximum likelihood estimator with a fitting radius of 5 pixels and an initial sigma of 1.6 pixels. STORM events were then filtered so that only events with the following criteria were retained: intensity >650 photons, sigma (range between 80 and 200) and uncertainty <20 nm. We did not perform sample drift correction but adjusted for re-blinking events by merging filtered events that were localised within 50 nm and 20 frames of the initial detection. Coordinate-based colocalisation was carried out using ThunderSTORM, with a step size of 50 nm for a total of 5 steps. CBC data was processed and analysed using GraphPad Prism and Microsoft Excel.

### SDS-PAGE, Phos-tag SDS-PAGE and western blotting

For normal SDS-PAGE, cells were lysed in RIPA buffer (R0278; Sigma Aldrich) supplemented with 1X protease inhibitor cocktail set 1 (539131; Sigma Aldrich). Lysates were diluted with 4X reducing LDS sample buffer (NP0007; Invitrogen) supplemented with 40 mM dithiothreitol (DTT; R0862; Thermo Fisher Scientific) to a 1X final concentration. Samples were loaded into 4-12% (w/v) NuPAGE Novex Bis-Tris gels (NP0321; Invitrogen) and Precision Plus Protein All Blue Standards (1610373: BioRAD) were used as molecular weight markers. Samples were separated by SDS-PAGE using NuPAGE MES SDS Running Buffer (NP0002; Invitrogen) at 150V for 45-60 mins. Proteins were then transferred onto a blotting membrane using the Trans-Blot Turbo Transfer System (1704150; Bio-Rad). The type of blotting membrane and length of transfer depended on the antibody added. Following transfer, the membrane was blocked using Casein Blocking Solution (37528; Sigma Aldrich) or 5% milk (LP0033B; Sigma Aldrich) diluted in Tris-buffered Saline, 0.1% (v/v) Tween 20 detergent (TBST; 655205; Thermo Fisher Scientific) for 1 h at RT or overnight at 4°C. Primary antibodies were added to the membrane for 1 h at RT or overnight at 4°C, followed by three 10 minute washes with TBST. Secondary antibodies were added to the membrane for 30-45 mins at RT, followed by three 10 mins washes with TBST. The membrane was then visualised using a LiCOR Odyssey scanner (LiCOR).

For Phos-tag SDS-PAGE, cells were lysed in RIPA buffer supplemented with 1X Halt™ Protease and Phosphatase inhibitor cocktail (78441; Thermo Fisher Scientific). Lysates were diluted with 3X reducing SDS sample buffer (200 mM Tris/HCl pH6.8 (BP152-1; Thermo Fisher Scientific), 30 mM DTT, 277 mM Sodium Dodecyl Sulfate (SDS; 436143; Sigma Aldrich), 6 mM Bromophenol Blue (B0126; Sigma Aldrich) and 30% (v/v) glycerol (G5516; Sigma Aldrich) to a 1X final concentration. Phos-tag SDS-PAGE was performed using custom-made gels following the manufacture’s protocol. Briefly, 20 μM Phos-tag affinity reagent (AAL-107; Wako Chemicals) was incorporated into 6% polyacrylamide gel containing MnCl_2_. Before transfer, the gel was washed twice for 10 mins in transfer buffer containing 10 mM EDTA, and then once with transfer buffer alone, using the Trans-Blot Turbo Transfer System reagents. The gel was transferred to a 0.45 μm PVDF membrane (GE10600119; Thermo Fisher Scientific) for 30 mins on the standard setting in the Trans-Blot Turbo Transfer System. The membrane was blocked in 5% milk TBST, and the antibodies were incubated in 1% milk TBST. Proteins were visualised by chemiluminescence using secondary antibodies conjugated with HRP and activated by Clarity Western ECL Substrate (1705060; Bio-Rad).

### MS-based proteomics

APEX2-expressing cells were treated with BP for 30 mins at 37°C, 5% CO_2_. If APEX2-expressing cells were co-cultured with Raji cells, the ligand-containing cells were added after 25 mins of incubation with BP at a ratio of Jurkat:Raji of 2:1, centrifuged at 50 g for 30 seconds and incubated at 37°C, CO_2_ for 5 mins. Raji cells were pre-pulsed with SEE using the protocol previously stated. Following the 30-minute incubation with BP, H_2_O_2_ was added for 2 mins at RT and immediately after incubation placed on ice. To halt proximity biotinylation the samples were washed three times with cold quenching solution (20 mM sodium ascorbate (PHR1279; Sigma Aldrich), 10 mM Troxol (238813; Sigma Aldrich) and 20 mM sodium azide (S2002; Sigma Aldrich) in DPBS). The cell pellet was resuspended in RIPA buffer supplemented with 1X protease inhibitor, mixed, and stored on ice for 20 mins. The lysate was centrifuged at maximum speed for 15 mins and the supernatant stored.

The concentration of cell lysates were quantified using Pierce 660nm (22660; Thermo Fisher Scientific), and this was used to determine the volume of sample required for addition to the MagReSyn® Streptavidin beads (MR-STV; ReSyn Biosciences). For every 100 μg of protein, 10 μL of MagReSyn® Streptavidin were used and approximately 800 μg of protein were enriched per sample. MagReSyn® Streptavidin beads were washed twice using RIPA lysis buffer and the prepared beads were incubated with the cell lysate overnight at 4°C with rotation. The beads were pelleted using a magnetic rack, and the supernatant removed. The beads were washed for 10 mins with rotation twice with wash buffer 1 (2% SDS), once with wash buffer 2 (0.1% deoxycholate (30970; Sigma Aldrich), 1% Triton X-100, 500 mM NaCl (S-3160-65; Thermo Fisher Scientific), 1 mM EDTA, and 50 mM HEPES pH 7.5 (BP310; Thermo Fisher Scientific)), and once with wash buffer 3 (0.5% NP-40 (85124; Thermo Fisher Scientific), 0.5% deoxycholate, 1 mM EDTA, and 10 mM Tris.HCl pH 7.4). After the final wash, the buffer was removed, and the material on the beads was eluted with 50 μL 5% SDS, 50 mM triethylammonium bicarbonate (TEAB; 17902; Fluka Analytical), 50 mM DTT, and 100 μM biotin (B4501; Sigma Aldrich) at 85°C for 10 mins with shaking at 1000 rpm.

Following elution, samples were prepared for mass spectrometry analysis using the S-Trap sample preparation (Profiti). Samples were alkylated in the dark for 30 mins with iodoacetamide (I6125; Sigma Aldrich) to a final concentration of 40 mM. Aqueous phosphoric acid (100573; Sigma Aldrich) at a final concentration of 1.2% in S-Trap binding buffer (90% aqueous methanol containing a final concentration of 100mM TEAB, pH 7.1) was added to the samples prior to loading onto the S-Trap columns (ProtiFi). After washes with S-Trap binding buffer, the samples were digested for 1 h at 47°C on-column with 1 μg sequencing grade modified trypsin (V5111; Promega) in digestion buffer (50 mM TEAB). Peptides were eluted from the column initially with 0.1% formic acid in water (85170; Thermo Fisher Scientific) and then with 0.1% formic acid in 30% acetonitrile (85175; Thermo Fisher Scientific). Eluted peptides were desalted using R3 beads (11339; Thermo Fisher Scientific) loaded onto Corning FiltreEX desalt filter plates (3505; Corning). Peptides were added to the filter plates with R3 beads, washed with 0.1 % formic acid in water, and eluted with 0.1 % formic acid in 30% acetonitrile. The desalted peptides were dried using a vacuum centrifuge and ran for 1 hr on the Q Exactive (Thermo Fisher Scientific) or Orbitrap Exloris 480 (Thermo Fisher Scientific) mass spectrometer.

Raw files were analysed using Proteome Discoverer (Thermo Fisher Scientific). Proteome Discoverer was used to search for proteins using the human UniProt database, applying the conditions specific to the mass spectrometers and modifications listed in Table 1. Further analysis of the raw abundances obtained from Proteome Discoverer was performed using Manchester Proteome Profiler. The mass spectrometry proteomics data have been deposited to the ProteomeXchange Consortium (http://proteomecentral.proteomexchange.org) via the PRIDE partner repository^43^ with the dataset identifier PXD063349.

**Table 1:** Proteome Discoverer search parameters.

| Tolerances |  |
| --- | --- |
| Precursor Mass Tolerance | 10 ppm |
| Fragment Mass Tolerance | 0.02 Da |
| Dynamic Modifications |  |
| Oxidation | +15.995 Da (M) |
| Biotin-tyramide | +361.146 Da (Y) |
| Acetyl | +42.011 Da (N-terminus) |
| Met-loss | -131.040 Da (M) |
| Met-loss+Acetyl | -89.030 Da (M) |
| Static Modifications |  |
| Carbamidomethyl | +57.021 Da (C) |

### Generation of liposomes and planar lipid bilayers

Planar lipid bilayers (PLB) were comprised of lipids obtained from Avanti Polar Lipids Inc. and were integrated into liposomes through an extrusion technique, as outlined in prior studies^44,45^. Bilayers were composed of lipid mixtures containing varying proportions of 4 mM stock solutions of DOPC (1,2-dioleoyl-sn-glycero-3-phosphocholine), Ni-NTA (1,2-dioleoyl-sn-glycero-3-[(N-(5-amino-1-carboxypentyl)iminodiacetic acid)succinyl]), and Cap-Biotin (1,2-dioleoyl-sn-glycero-3-phosphoethanolamine-N-(CAP biotinyl)). To create PLBs, 8-well bottomless sticky glass slides (80828; Ibidi) were mounted onto cleanroom cleaned coverslips (1472315; NEXTERION®; SCHOTT), securely fastened using a clamp (80040; Ibidi), and incubated for 8 hr at 37°C to adhere. A liposome solution containing 50% Ni-NTA, 45% DOPC, and 5% Cap-Biotin in 200 μL were added to each well. Following a 20 mins incubation at RT, the wells were washed 5 times by sequential addition and removal of DPBS to remove excess lipid. 1% BSA with 100 μM nickel sulphate was added to block for 1 hr at RT. 1 μg/mL streptavidin was added and incubated for 20 mins at RT, and excess streptavidin was removed by 5 times DPBS washes. Ligands were loaded at 400 mol/μm^2^ (0.5 μg/mL) for 1 h at 37°C, 5% CO_2_. All conditions included His-ICAM-1 (Peak Proteins), and depending on the condition, also included His-CD155 (2530; R&D Systems), His-CD111 (AVI2880; R&D Systems), and monobiotinylated OKT3 (AB467057; Thermo Fisher Scientific). The ligands were washed 5 times in DPBS, and 100,000 cells were added onto the ligand-loaded PLBs for 5 mins at 37°C, 5% CO_2_ and fixed using a final concentration of 4% PFA for 20 mins at 37°C, 5% CO_2_. The wells were washed 5 times in DPBS, permeabilised with 0.1% Triton X-100 for 10 mins at RT, and washed again. The wells were blocked with 3% BSA/1% HS for 1 hr at RT, stained with antibodies at 5 μg/mL for 1 hr at RT, and washed before imaging by TIRF microscopy.

### Enzyme-linked immunosorbent assay (ELISA)

IL-2 levels in cell supernatants were quantified by an enzyme-linked immunosorbent assay (ELISA). Nunc MaxiSorp 96-well plates (44-2404-21; Thermo Fisher Scientific) were coated overnight at 4°C with 1 μg/mL human IL-2 capture antibody (555051; BD Pharmingen) in 50 mM carbonate bicarbonate buffer (C3041; Sigma Aldrich). After three washes with wash buffer (DPBS with 0.05% Tween20 (P1379: Sigma Aldrich) to remove excess antibody, wells were blocked with blocking buffer (1% BSA in DPBS, 0.22 μm filtered) for 1 hr at RT. Samples and standards (rhIL-2; 554603; BD Biosciences) were added to the coated wells, incubated for 2 hrs at RT, and washed 3 times. 1 μg/mL detection antibody (555040; BD Pharmingen) was added, incubated for 1 hr at RT, and washed 3 times. Streptavidin-HRP (554066; BD Biosciences) was applied for 30 mins at RT, followed by 3 washes. TMB solution (54827-17-7; MP Bio) was then added, allowing the reaction to proceed for 10 mins, after which it was stopped with 0.5 M H_2_SO_4_ (339741; Sigma Aldrich). Absorbance was measured at 450 nm on a spectrophotometer.

### Quantification and Statistical Analysis

Statistical analyses were carried out using GraphPad Prism software. To determine whether the data followed a normal distribution, the Shapiro-Wilk test was applied to each dataset. For comparing two groups with normally distributed data, a two-tailed Student’s t-test was employed. If one or both groups did not meet the normality assumption, a Mann-Whitney test was used. For comparisons involving three or more groups, statistical significance was evaluated using one-way or two-way ANOVA. Non-significant differences were defined as p ≥ 0.05, while statistically significant differences were denoted as * p<0.05, ** p<0.01, *** p<0.001, and **** p<0.0001. Datasets were presented as mean ± standard deviation (SD).

## Acknowledgements

The authors thank Hayley Bennett and Anthony Adamson for generation of the APEX2 constructs, Gareth Howell and Bradley Dean for help with fluorescence-assisted cell sorting, and Stacey Warwood and Emma-Jayne Keevill in the Bio-MS facility for help with mass spectrometry analysis. The authors would also like to thank all members of the Worboys, Davis and Humphries labs for supportive discussions.

This work was supported by a Wellcome Career Development Award (307027/Z/23/Z to J.D.W), a Wellcome Investigator Award (110091/Z/15/Z to D.M.D.), a Medical Research Council Research Grant (MR/W031698/1 to D.M.D.), a Cancer Research UK Programme Grant (DRCRPG-100002 to M.J.H.), and a PhD studentship funded by Wellcome (to W.H.Z). The work was also conducted within the Wellcome Centre for Cell-Matrix Research (core award 203128/Z/16/Z).

## Author contributions

Conceptualization, W.H.Z., M.J.H., D.M.D. and J.D.W.; Data curation, W.H.Z.;

Formal Analysis, W.H.Z,, S.A.C. and J.D.W.; Funding acquisition, M.J.H., D.M.D. and J.D.W.; Investigation, W.H.Z. and J.D.W.;

Methodology, W.H.Z. and J.D.W.; Supervision, M.J.H., D.M.D. and J.D.W.; Visualization, W.H.Z. and J.D.W.;

Writing – original draft, W.H.Z. and J.D.W.;

Writing – review & editing, W.H.Z., M.J.H., D.M.D. and J.D.W.;

