## Supplementary information for "Inhibitory TIGIT signalling is dependent on T cell receptor activation"

### TIGIT phosphorylation and signalling in T cells is dependent on both ligation and T cell activation

Supplementary figures

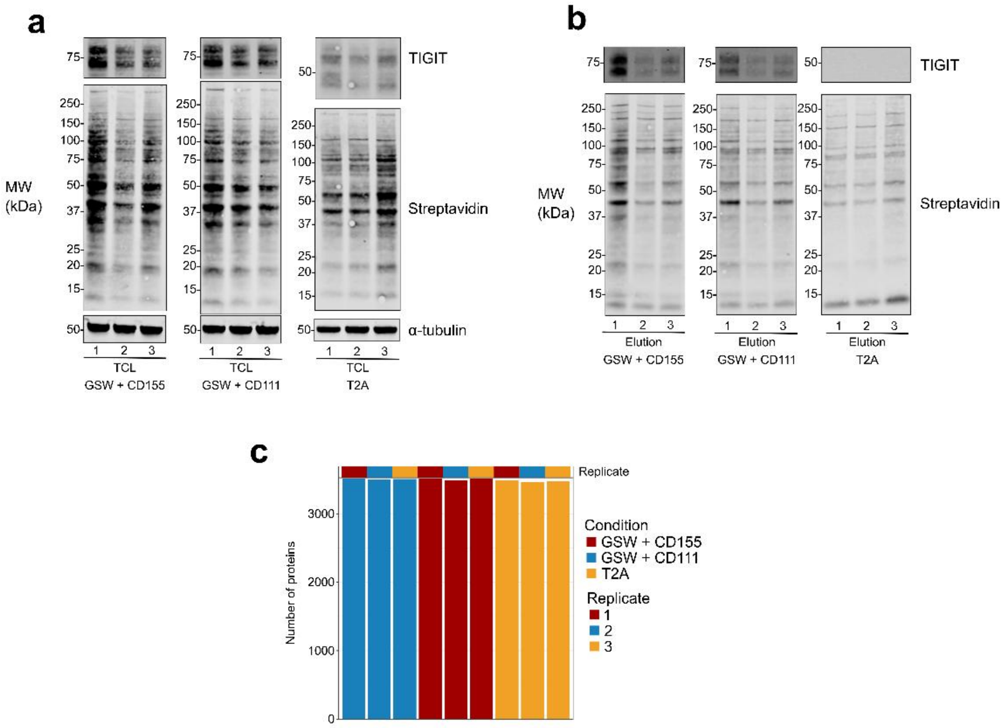


**Supplementary Figure 1 – Western blotting of samples used for ligation-dependent proximity proteomics and additional data from LC-MS/MS analysis. a.** Western blots showing TIGIT and biotinylated proteins in TIGIT-GSW-APEX2 (GSW) Jurkat cells co-cultured with either Raji-CD111 or Raji-CD155 in the presence of superantigen, alongside the background control Jurkat cells expressing TIGIT-T2A-APEX2 (T2A). We show the banding patterns for the total cells lysates (TCL) from three replicates of each condition following proximity labelling. A loading control blot for α-Tubulin (DM1A) is also shown below from the same lysates. Molecular weight markers are depicted on the left. **b.** Western blots showing TIGIT and biotinylated proteins in streptavidin enriched eluates from the TCL in **a**. Molecular weight markers are depicted on the left **c.** Total number of proteins identified by LC-MS/MS analysis of the eluates in **b**.

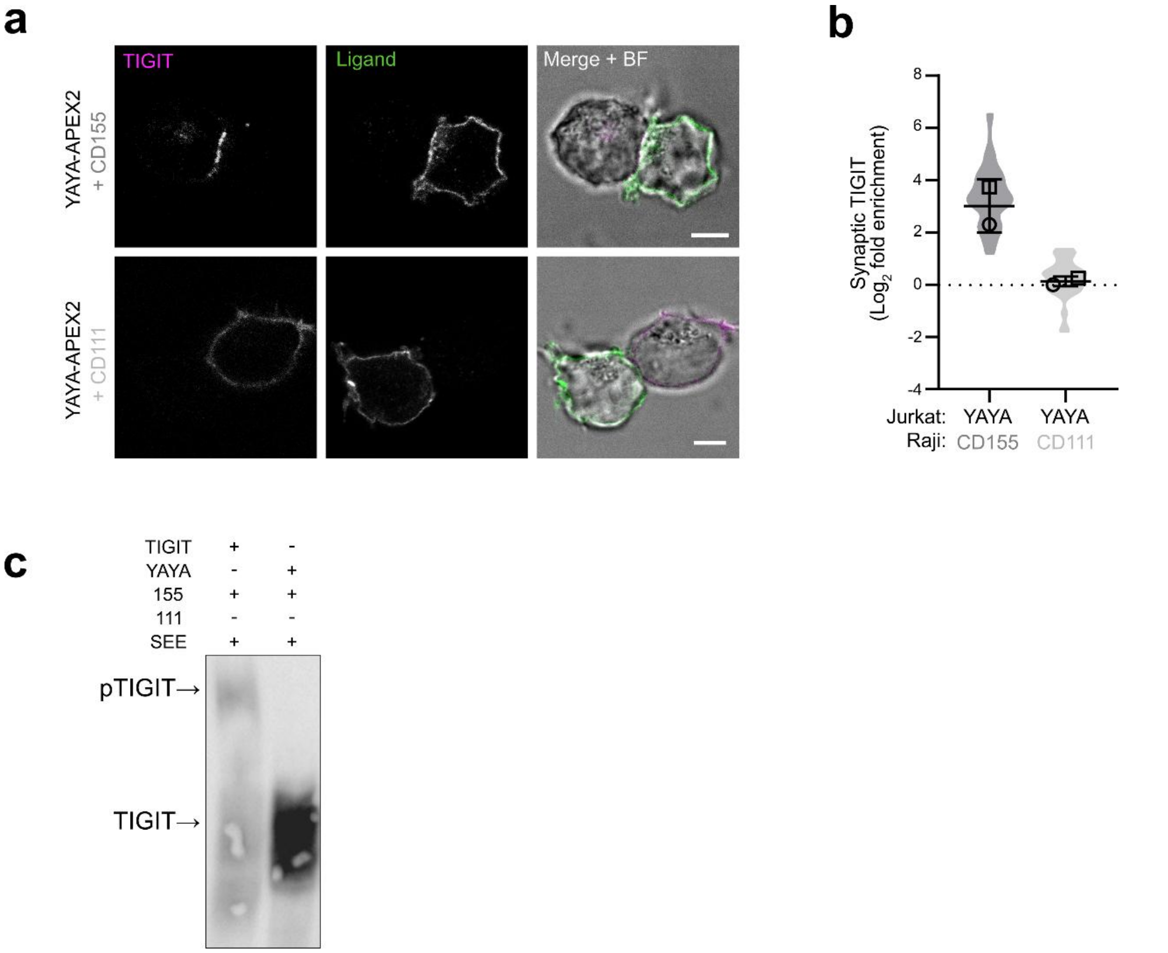


**Supplementary Figure 2 – Generation of a functional mutant TIGIT-APEX2 fusion protein to perform proximity proteomics in T cells. a.** Confocal microscopy images showing TIGIT (magenta) in signalling deficient TIGIT-GSW-APEX2 (YAYA) Jurkat cells that were conjugated with different Raji cell populations (expressing either CD111 or CD155) for 5 mins, as indicated on the left of the panel. The V5 stain labels expressed ligands on Raji cells (green). A merged fluorescence-BF image is provided. Scale bars = 5 µm. **c.** Mean $\log_2$ fold change in synaptic TIGIT enrichment in Jurkat cells, from the conjugates shown in **a** (±S.D.; n=2 independent experiments). **c.** Western blot analysis of TIGIT using either Phos-tag SDS-PAGE (top) or standard SDS-PAGE (middle) to examine TIGIT phosphorylation in YAYA Jurkat cells conjugated to SEE-pulsed Raji CD155 or CD111 cells for 5 mins, as indicated.

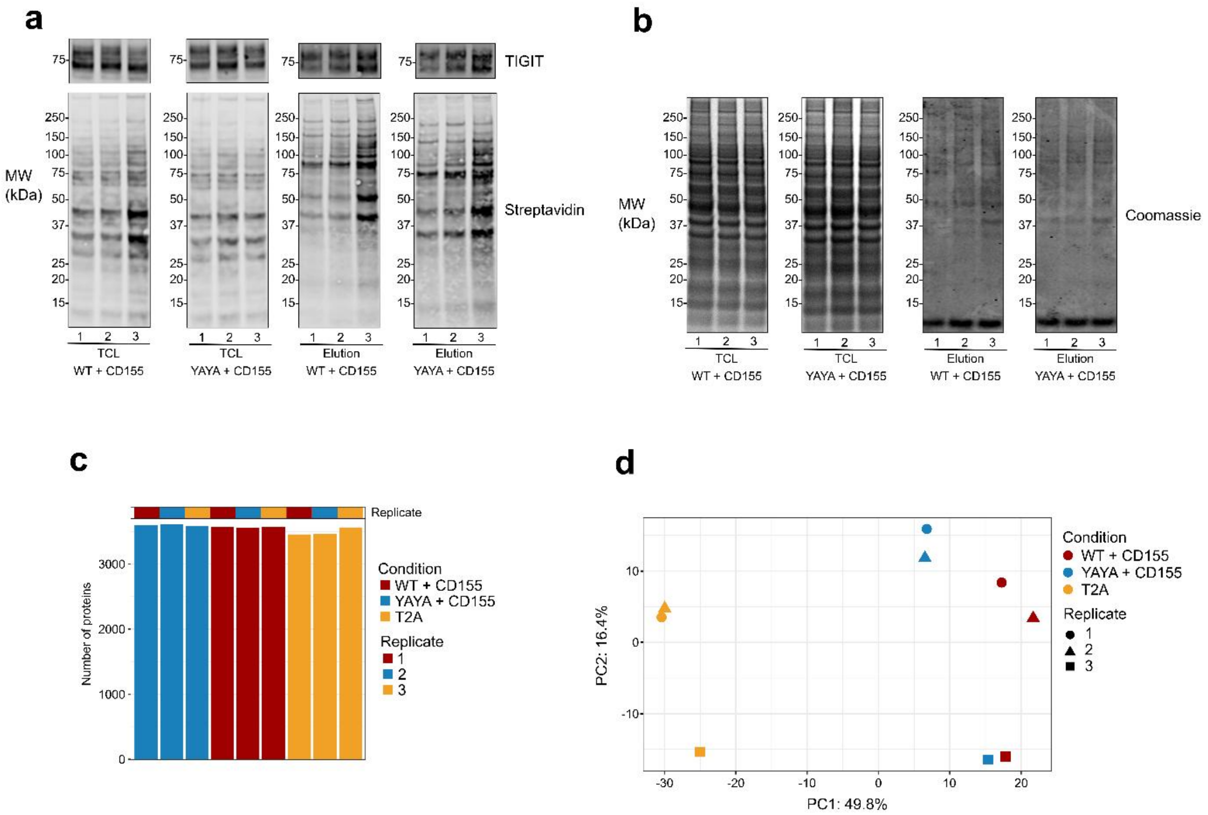


**Supplementary Figure 3 – Western blotting of samples used for inhibitory motif-dependent proximity proteomics and additional data from LC-MS/MS analysis. a.** Western blots showing TIGIT and biotinylated proteins in TIGIT-GSW-APEX2 (WT) and the signalling-deficient form, YAYA-GSW-APEX2 (YAYA), Jurkat cells co-cultured with Raji-CD155 in the presence of superantigen, alongside the background control Jurkat cells expressing TIGIT-T2A-APEX2 (T2A). We show the banding patterns for the total cells lysates (TCL) from three replicates of each condition following proximity labelling and the streptavidin enriched eluates from the TCL. Molecular weight markers are depicted on the left. **b.** Total protein control (Coomassie) from the same TCL and eluates in **a**. **c.** Total number of proteins identified by LC-MS/MS analysis of the eluates in **a**. Principal component analysis depicting the relative positions of each replicate of the conditions analysed according to the variation in the data, with principal components 1 (PC1) and 2 (PC2) plotted.

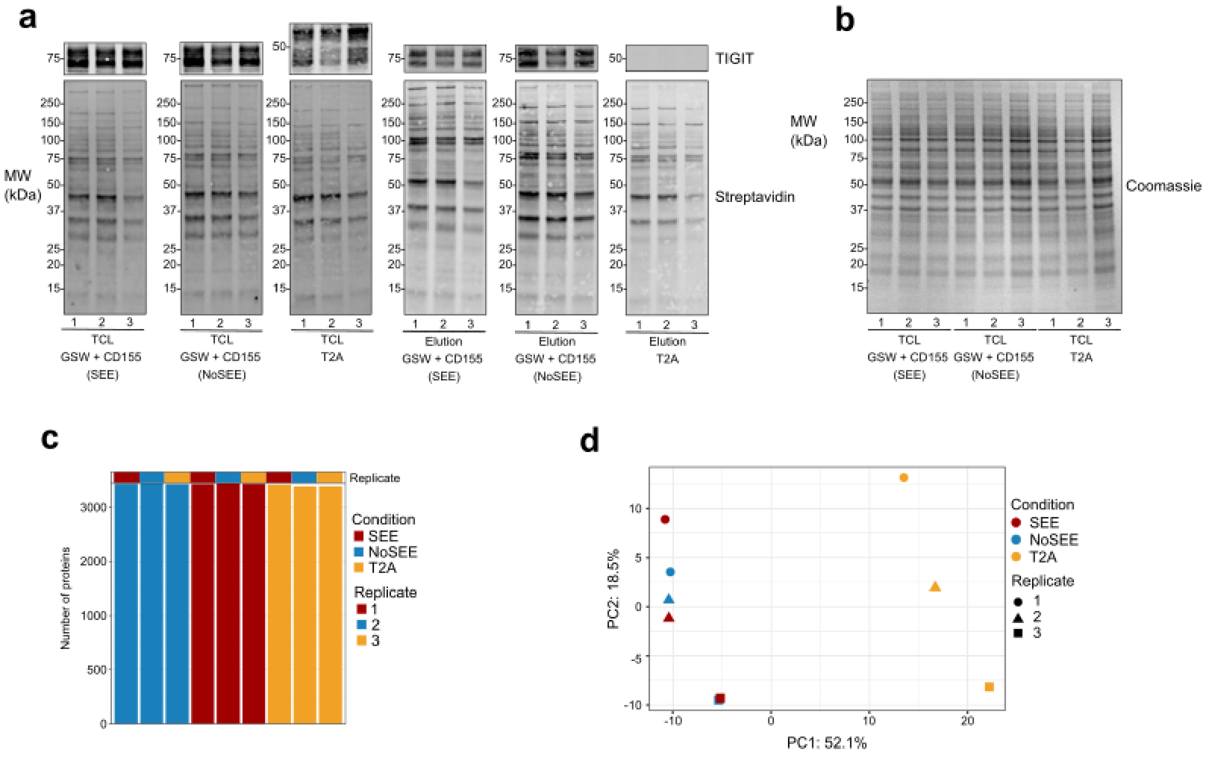


**Supplementary Figure 4 – Western blotting of samples used for TCR activation-dependent proximity proteomics and additional data from LC-MS/MS analysis. a.** Western blots showing TIGIT and biotinylated proteins in TIGIT-GSW-APEX2 Jurkat cells co-cultured with Raji-CD155 with or without SEE-pulsing, alongside the background control Jurkat cells expressing TIGIT-T2A-APEX2 (T2A). We show the banding patterns for the total cells lysates (TCL) from three replicates of each condition following proximity labelling and the streptavidin enriched eluates from the TCL. Molecular weight markers are depicted on the left. **b.** Total protein control (Coomassie) from the same TCL and eluates in **a**. **c.** Total number of proteins identified by LC-MS/MS analysis of the eluates in **a**. Principal component analysis depicting the relative positions of each replicate of the conditions analysed according to the variation in the data, with principal components 1 (PC1) and 2 (PC2) plotted.

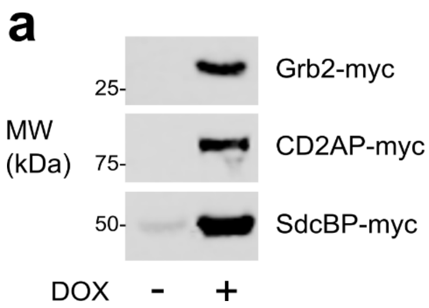


**Supplementary Figure 5 – Western blot analysis of protein expression in TIGIT-SNAP Jurkat cells**

**a.** Western blot analysis of myc-tagged proteins in TIGIT-SNAP expressing Jurkat cells stably expressing doxycycline inducible Grb2-, CD2AP-, or SdcBP-myc were generated using lentivirus. The cells were incubated with or without doxycycline for two days and then lysed. Lysates were analysed using an anti-myc antibody.

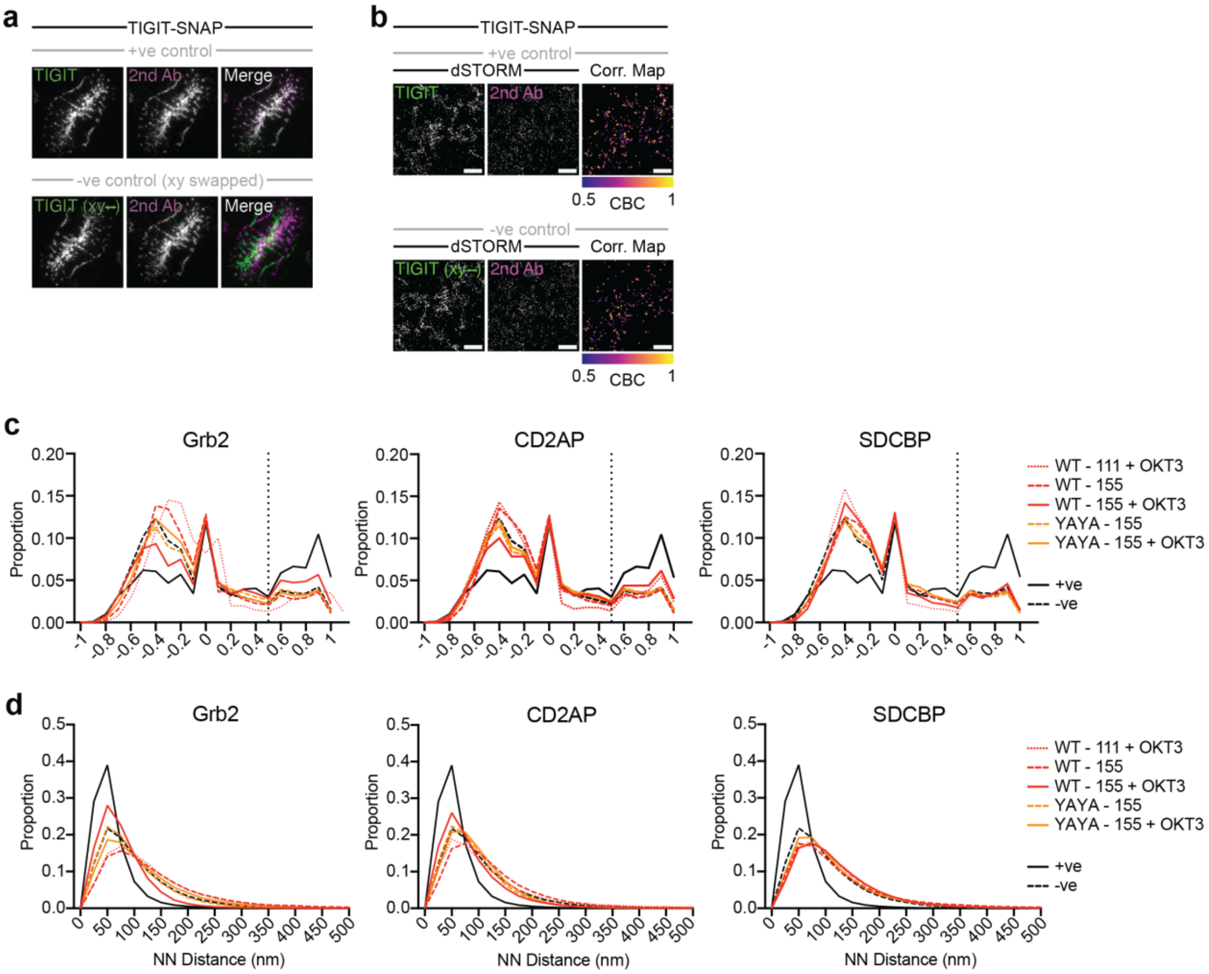


**Supplementary Figure 6 – Additional dSTORM analysis. a.** Representative Total Internal Reflection Fluorescence (TIRF) microscopy images of WT TIGIT-SNAP stained with a primary TIGIT antibody (green) and a secondary antibody (magenta) at the immune synapse of Jurkat TIGIT-SNAP cells that have interacted with planar lipid bilayers (PLB) loaded with ICAM-1 (100 molecules/μm$^2$), CD155 (400 molecules/μm$^2$), and OKT3 (100 molecules/μm$^2$) for 5 mins (top row; positive control). We invert the XY coordinates of the secondary antibody channel to create a suitable negative control (bottom row). A merged fluorescence image is also provided for each comparison, scale bars = 5 μm. **b.** TIRF dSTORM images of WT TIGIT-SNAP from the cells displayed in **a**. Each image represents a smaller 5 μm$^2$ region of the indicated TIRF image. For each condition, we also provide a map of TIGIT localisations (from primary antibody channel) where coordinate based colocalisation (CBC) values are ≥0.5, and colour code the localisations based on the exact CBC values (Correlation Maps). **c,d.** Histograms displaying the distribution of CBC values (**c**) and nearest neighbour distances (**d**) of TIGIT localisations compared to each of the indicated proteins, in the indicated conditions.

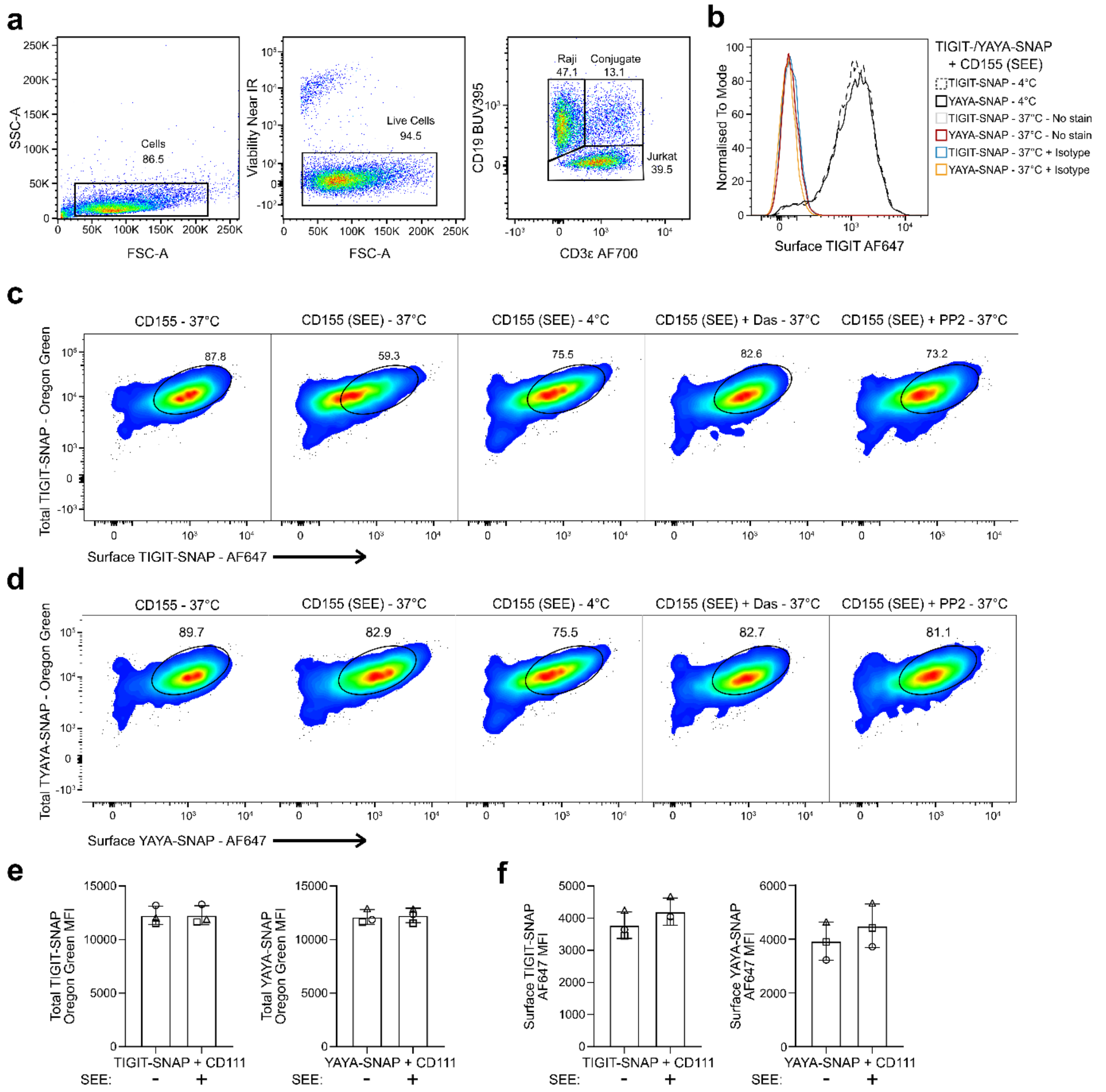


**Supplementary Figure 7 – Flow cytometry analysis of cell surface TIGIT. a.** Strategy used to gate on Jurkat and Raji cells. **b.** Histogram of cell surface TIGIT expression on TIGIT-SNAP/YAYA-SNAP Jurkat cells co-cultured with Raji-CD155 cells at 37°C or 4°C with or without SEE. Isotype and FMO staining controls with TIGIT-AF647 staining are shown. **c.** Flow cytometry analysis of TIGIT-SNAP Jurkat cells showing total TIGIT expression (SNAP-TAG) and cell surface TIGIT expression (antibody labelling) following indicated Raji co-culture conditions. **d.** Flow cytometry analysis of YAYA-SNAP Jurkat cells showing total TIGIT expression (SNAP-TAG) and cell surface TIGIT expression (antibody labelling) following indicated Raji co-culture conditions. **e.** Median Fluorescent Intensity values of total TIGIT expression when TIGIT-SNAP/YAYA-SNAP cells were co-cultured with Raji cells expressing CD111 control ligand. **f.** Median Fluorescent Intensity values of total TIGIT expression when TIGIT-SNAP/YAYA-SNAP cells were co-cultured with Raji cells expressing CD111 control ligand.
